# Hepatocyte-specific CLSTN3B ablation impairs lipid droplet maturation and alleviates diet-induced steatohepatitis in mice

**DOI:** 10.1101/2025.05.20.655199

**Authors:** Chuanhai Zhang, Dengbao Yang, Hiroyuki Suzuki, Jingxuan Chen, Jingjing Wang, Mengchen Ye, Jin Zhou, Qiyu Zeng, Meijuan Bai, Mei-Jung Lin, Jeon Lee, Hao Zhu, Yujin Hoshida, Xing Zeng

## Abstract

Excessive lipid accumulation in hepatocytes, a hallmark of metabolic dysfunction-associated steatotic liver disease (MASLD), can lead to progressive liver damage. Understanding the molecular mechanisms governing lipid storage in hepatocytes is essential for identifying therapeutic targets to halt MASLD progression. Here, we show a pivotal role for the protein calsyntenin 3β (CLSTN3B) in promoting lipid droplet (LD) maturation and lipid storage in hepatocytes. Previously characterized as an endoplasmic reticulum (ER)-LD contact protein that facilitates LD maturation in adipocytes, we now show that CLSTN3B expression is strongly induced in mouse hepatocytes by peroxisome proliferator-activated receptor gamma (PPARγ) in response to dietary caloric excess. Hepatocyte-specific deletion of CLSTN3B in mice significantly increases energy expenditure, reduces metabolic efficiency, and protects against diet-induced hepatic steatosis and fibrosis. Mechanistically, CLSTN3B deficiency causes reduced LD phospholipid coverage and increased lipase recruitment. This results in enhanced fatty acid oxidation driven by a futile cycle of lipolysis and re-esterification. Notably, human clinical data reveal a positive correlation between hepatic CLSTN3B expression and MASLD severity and progression, emphasizing its relevance to human disease. Together, our findings establish CLSTN3B as a key regulator of hepatocyte lipid storage and metabolic efficiency and highlight its potential as a therapeutic target in MASLD.

## Introduction

Metabolic dysfunction-associated steatotic liver disease (MASLD), formerly known as non-alcoholic fatty liver disease (NAFLD), is now recognized as the most prevalent chronic liver disorder worldwide, affecting more than 25% of the global population^1,2^. MASLD encompasses a spectrum of liver pathology that begins with the accumulation of excess neutral lipids, mostly triglycerides, in hepatocytes (steatosis) and can progress to inflammatory steatohepatitis, fibrosis, cirrhosis, and ultimately hepatocellular carcinoma^3,4^.

While steatosis itself is considered relatively benign, the transition to more advanced stages of liver damage is clinically significant and remains incompletely understood^5^. A key event in this progression is the activation of hepatic stellate cells (HSCs), which, in their quiescent state, store vitamin A as retinyl esters. Chronic lipid overload and oxidative stress cause hepatocyte damage, creating a necro-inflammatory microenvironment that induces HSCs to transdifferentiate into myofibroblast-like cells. These cells produce extracellular matrix components, including type I collagen, thereby driving liver fibrosis^5^.

Since excessive lipid deposition in hepatocytes is the initiating event in MASLD, therapeutic strategies have focused on reducing hepatic lipid burden and enhancing mitochondrial fatty acid oxidation (FAO). One promising avenue involves liver-targeted mitochondrial uncouplers, which dissipate the mitochondrial proton gradient, reduce metabolic efficiency, and drive increased substrate oxidation^6–10^. While these agents have shown efficacy in reducing steatosis in preclinical models, their narrow therapeutic window and risk of systemic toxicity (e.g., hyperthermia, energy depletion) have limited clinical translation^11^. Alternatively, the thyroid hormone receptor-β (THRβ) agonist resmetirom, recently approved by the U.S. Food and Drug Administration for MASLD patients with moderate to advanced fibrosis, modulates lipid metabolism transcriptionally, enhancing lipolysis, lipophagy, and mitochondrial oxidation of liberated fatty acids^12^.

A hallmark of hepatic steatosis is the emergence of numerous lipid droplets (LDs) in hepatocytes, dynamic organelles that store excess lipids primarily as triglycerides. LDs are bounded by a phospholipid monolayer and originates from the cytoplasmic face of the endoplasmic reticulum (ER)^13^. During subsequent expansion, LDs continue to receive phospholipids and proteins from the ER^14^. Cytoplasmic proteins are later recruited to the LD surface, often through amphipathic motifs that recognize packing defects in the phospholipid monolayer, to promote LD maturation^15,16^. This surface, composed of a phospholipid monolayer and associated proteins, serves to restrict lipase access to the neutral lipid core, thereby facilitating lipid storage and protecting cells from lipotoxicity by sequestering bioactive fatty acids and derivatives in inert, neutral forms^17–19^. While this compartmentalization can be protective, it may also contribute to the progression of MASLD by enabling chronic lipid accumulation. Supporting this view, human genetic studies have identified key roles for LD-associated proteins in MASLD pathogenesis. For example, the I148M variant in patatin-like phospholipase domain-containing pro-tein 3 (PNPLA3), one of the strongest genetic risk factors for MASLD, impairs triglyceride hydrolysis by sequestering α/β hydrolase domain-containing protein 5 (ABHD5, also known as CGI-58), thereby inhibiting adipose triglyceride lipase (ATGL)-mediated lipolysis^20–24^. Conversely, loss-of-function mutations in the cell death-inducing DFFA-like effector protein B (CIDEB) protein, a protein localizing to ER and LD, confer protection against liver disease^25,26^. Genetic or pharmacological perturbation of CIDEB improves MASLD condition in mice by markedly reducing LD volume and enhancing FAO^27–30^. Similarly, silencing perilipin 2 (PLIN2), a major LD-coating protein, or diacylglycerol O-acyltransferase 2 (DGAT2), the enzyme responsible for triglyceride synthesis and driving LD biogenesis, both mitigates MASLD phenotypes^31–35^. Together, these findings suggest that modulating LD biogenesis and surface composition may offer a promising therapeutic approach to counteract pathological lipid storage in hepatocytes.

Recent studies in adipocytes identified calsyntenin 3β (CLSTN3B) as a novel protein localizing to ER-LD contact sites via an N-terminal LD-targeting domain and a C-terminal ER transmembrane region^36,37^. It promotes LD expansion and maturation by facilitating phospholipid transfer from the ER to the LD surface through a unique arginine-rich motif that mediates hemifusion-like membrane interactions^36^. In adipocytes, loss of CLSTN3B reduces LD phospholipid coverage, increases basal lipolysis, and limits lipid storage capacity^36^. Under normal conditions, CLSTN3B expression is largely restricted to adipocytes^38^, likely reflecting its role in supporting the maturation of uniquely large LDs characteristic of adipocytes. However, hepatocytes in MASLD also acquire large LDs, raising the possibility that CLSTN3B may play a previously unrecognized role in hepatic lipid storage and disease progression.

Here, we show that dietary caloric excess induces hepatic CLSTN3B expression through peroxisome proliferator-activated receptor gamma (PPARγ) activation, driving LD expansion and lipid accumulation in hepatocytes. Hepatocyte-specific deletion of CLSTN3B impairs LD surface phospholipid coverage, facilitates lipase access, and promotes futile lipolysis/re-esterification cycling, thereby enhancing FAO and energy expenditure. This metabolic rewiring reduces hepatic steatosis, inflammation, and fibrosis, all of which are exacerbated by CLSTN3B overexpression. Furthermore, CLSTN3B expression in MASLD patient liver samples positively correlates with fibrosis stage and disease progression. Collectively, our findings uncover CLSTN3B as a key regulator of hepatocellular LD biogenesis and a driver of MASLD pathogenesis, offering new insights into the ER-LD interface as a therapeutic target.

## Results

### Dietary caloric overload induces CLSTN3B expression in mouse hepatocytes

To promote LD formation and lipid storage in hepatocytes, we subjected mice to one of the three fat-enriched diets: a high-fat diet (HFD, 60% fat), a western diet (WD, 21.1% fat, 41% sucrose, 1.25% cholesterol by weight) supplemented with a high-sugar solution (23.1 g/L D-fructose and 18.9 g/L D-glucose), and a choline-deficient high fat-diet (CDAHFD, 60% fat). We then analyzed *Clstn3b* expression in the liver. Hepatic *Clstn3b* mRNA level increased markedly over time in response to the dietary interventions (a 40-50 fold induction after 16 weeks of HFD or WD, and a 12-fold induction at 12 weeks of CDAHFD, Fig. 1a). To determine whether the CLSTN3B protein is similarly upregulated, we isolated hepatocyte LDs from mice maintained on those diets for 12 weeks and analyzed CLSTN3B expression by Western blot. Consistent with previous observations in adipocytes^36^, full-length CLSTN3B, and to a lesser extent, a C-terminal fragment, was detected in LDs from mice fed with the high-fat diets, but not from chow-fed controls (Fig. 1b). Both CLSTN3B bands were absent in hepatocyte LDs isolated from the liver-specific *Clstn3b* knockout (KO) mice (Fig. 1c) (LKO: *albumin-cre*; *Clstn3b^fl/fl^*) on HFD. Fluorescence microscopy revealed that endogenous CLSTN3B localizes to the LD surface in hepatocytes isolated from wild-type (WT) control but not the LKO mice (Fig. 1d-e).

**Figure 1.**
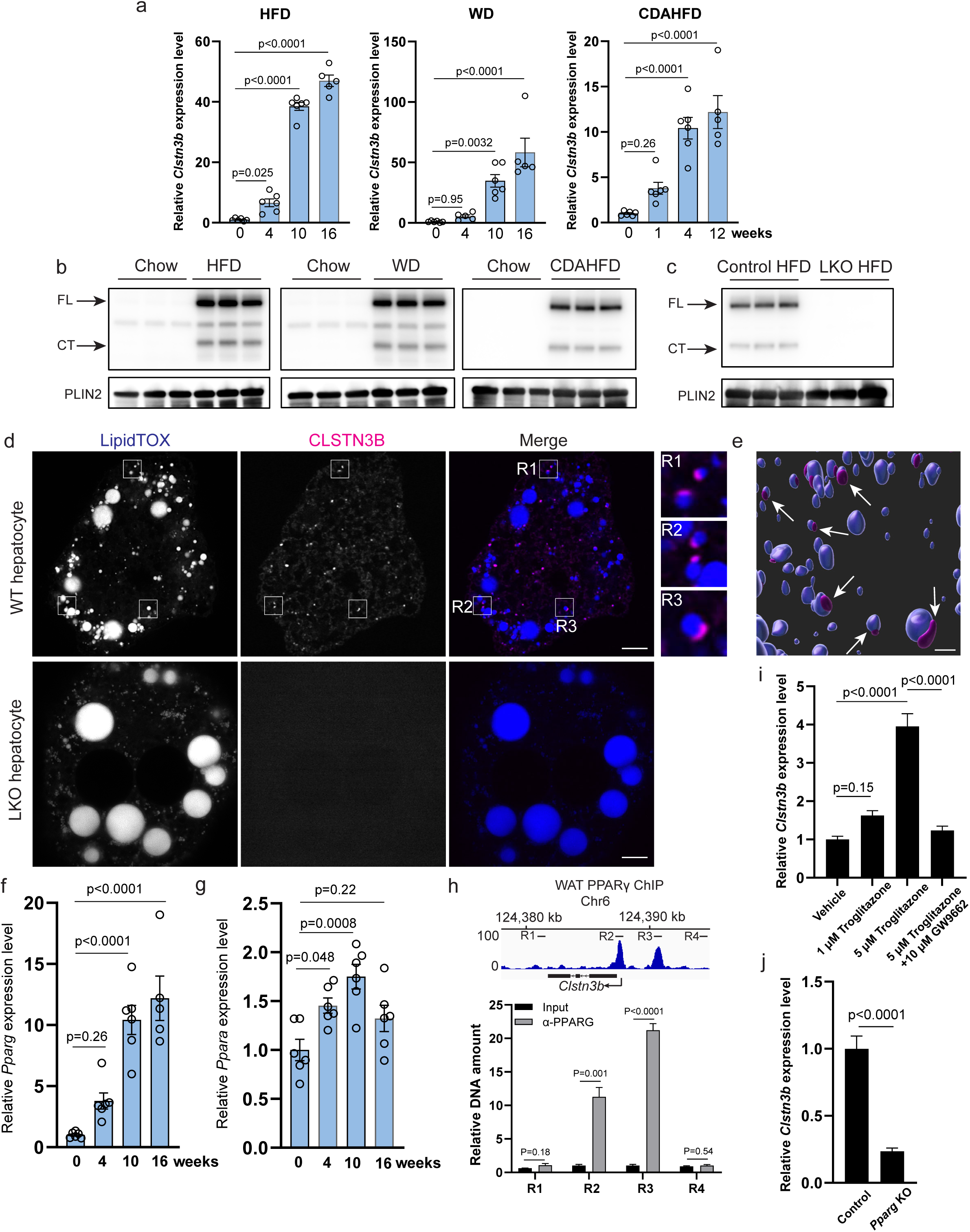
Dietary caloric overload induces CLSTN3B expression in mouse hepatocytes. **a**, qPCR analysis of liver *Clstn3b* expression from mice on different diets (nL=L6). **b**, Western blot analysis of CLSTN3B expression on LDs isolated from mice on different diets. **c**, Western blot analysis of CLSTN3B expression on LDs isolated from control or LKO mice on HFD. **d**-**e**, Fluorescence microscopic images (**d**) and 3D reconstruction (**e**) of CLSTN3B localization in hepatocytes. Scale bar: 5 μm in **d** and 1 μm in **e**. **f**-**g**, qPCR of liver *Pparg* (**f**) and *Ppara* (**g**) expression from mice on HFD (nL=L6). **h**, ChIP-qPCR analysis of PPARγ bindings sites upstream of the *Clstn3b* gene (nL=L3). PPARγ ChIP-seq at the same locus from the WAT is shown as a reference. **i**, effect of PPARγ ligands on *Clstn3b* expression in hepatocytes (nL=L4). **j**, qPCR analysis of liver *Clstn3b* expression from control or *Pparg* KO mice on HFD (nL=L6). Data are mean ± s.e.m. Statistical significance was calculated by One-way ANOVA with Tukey’s post hoc test in **a**, **f**, **g**, and **i**, and unpaired Student’s two-sided t-test in **h** and **j**.

Previous studies have shown that high-fat feeding induces expression of PPARγ, a nuclear receptor critical for adipogenesis, in hepatocytes, where it transcriptionally activates adipocyte-selective genes^39–41^. We found that *pparg* expression in the liver is induced in mice on HFD following a similar time course to *Clstn3b* (Fig. 1f). A recent report showed that hepatic *Clstn3b* expression is induced ∼3-fold upon fasting in a peroxisome proliferator-activated receptor alpha (PPARα)-dependent manner^42^. Compared to *Pparg*, *Ppara* transcript was only modestly induced by short-term HFD feeding and the difference from baseline became insignificant by 16 weeks (Fig. 1g), suggesting that PPARγ may be the dominant regulator of HFD-induced expression of *Clstn3b* in hepatocytes. To test this, we performed PPARγ ChIP-qPCR from hepatocytes and identified PPARγ binding sites upstream of the *Clstn3b* transcription start site positioned similarly to those white adipocytes (Fig. 1h). Treatment of hepatocytes with troglizatone, a PPARγ full agonist, significantly induced *Clstn3b* mRNA level in hepatocytes, an effect abolished by co-treatment with GW9662, a covalent non-agonistic PPARγ ligand (Fig. 1i). Finally, we induced liver-specific ablation of *Pparg* in mice on HFD by delivering AAV8-sgRNA targeting *Pparg* into CAS9 expressing mice after 2 weeks of HFD feeding. This resulted in a 72% reduction of hepatic PPARγ protein levels and an ∼80% reduction of *Clstn3b* transcript levels (Fig. 1j, Fig. S1a-c). Taken together, these findings establish PPARγ as a key transcriptional driver of *Clstn3b* expression in hepatocytes upon dietary caloric excess.

### Liver specific deficiency of CLSTN3B alleviates HFD-induced hepatic steatosis

To investigate the physiological significance of CLSTN3B in the liver, we analyzed male LKO mice and WT control littermates after 10 weeks of HFD feeding, starting at 6-8 weeks of age. Compared to WT controls, the LKO mice displayed significantly reduced liver mass, liver-to-body weight ratio, and liver triglyceride content (Fig. 2a-c). A modest, non-significant trend toward reduced body weight was noted in LKO mice (Fig. 2d, p=0.106). No concomitant increases in serum cholesterol, triglyceride and free fatty acid (FFA) levels, as well as adipose tissue mass were observed (Fig. 2a, e-f), suggesting that the reduction in hepatic lipid accumulation was not due to enhanced lipid export or redistribution to peripheral tissues. LKO mice also exhibited significantly lower serum aspartate aminotransferase (AST) and alanine aminotransferase (ALT) levels (Fig. 2g), consistent with alleviated liver injury. Gross liver appearance and histological analysis further confirmed markedly reduced lipid accumulation in LKO livers (Fig. 2h). Interestingly, in contrast to numerous small LDs accumulated in WT hepatocytes, CLSTN3B-deficient hepatocytes tend to harbor a small number of substantially larger LDs (10-20 μm in diameter, see the inset of Fig. 2h), indicative of abnormal LD biogenesis.

**Figure 2.**
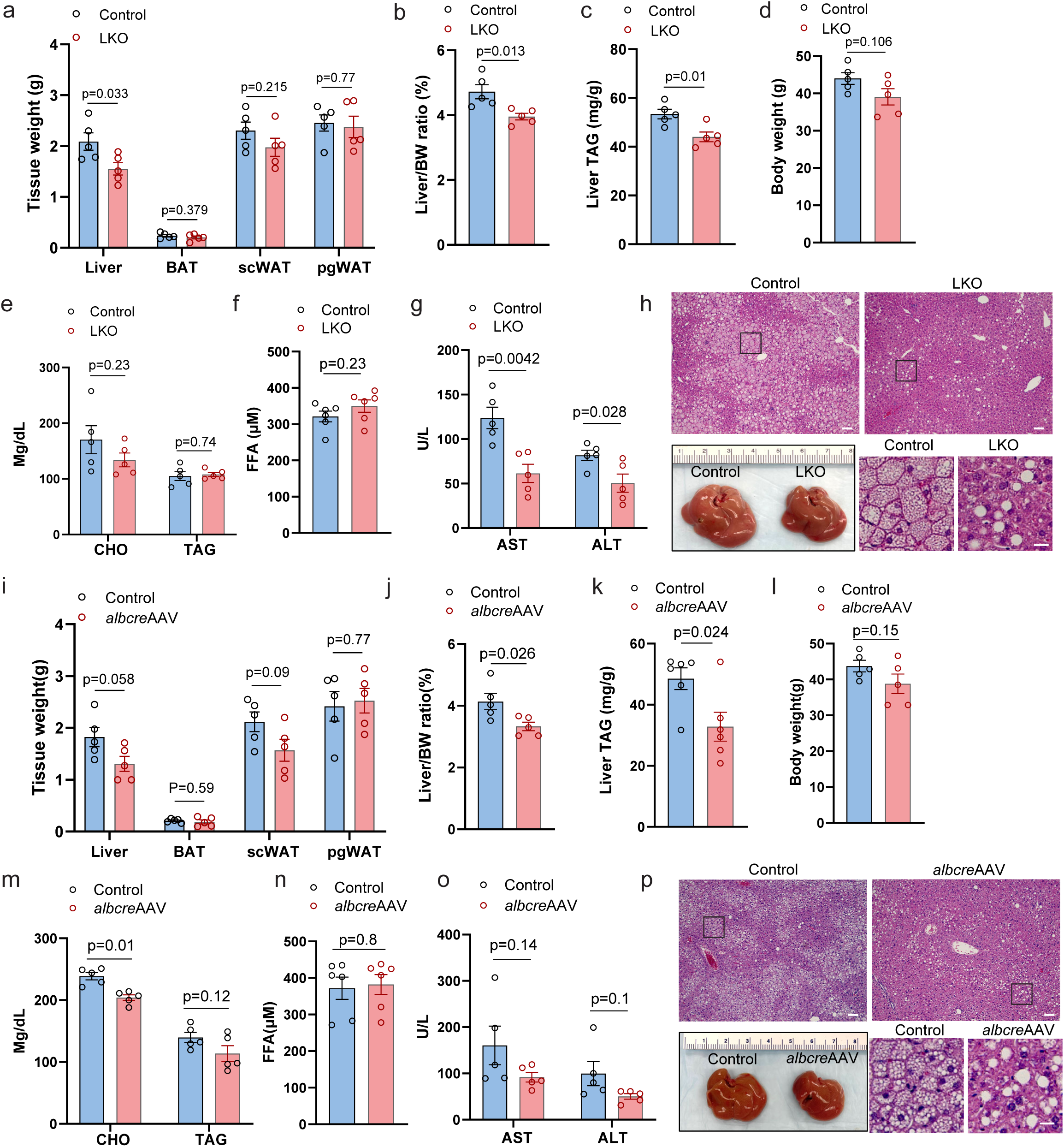
Liver specific deficiency of CLSTN3B alleviates HFD-induced hepatic steatosis. **a**-**h**, Tissue weight (**a**), liver/body weight ratio (**b**), liver TAG content (**c**), body weight (**d**), serum cholesterol and TAG level (**e**), serum FFA level (**f**), serum AST and ALT level (**g**), liver gross appearance and histology (**h**) from WT control and LKO mice on HFD for 10 weeks (n=5). **i**-**p**, Tissue weight (**i**), liver/body weight ratio (**j**), liver TAG content (**k**), body weight (**l**), serum cholesterol and TAG level (**m**), serum FFA level (**n**), serum AST and ALT level (**o**), liver gross appearance and histology (**p**) from *Clstn3b^fl/fl^* mice receiving control or *alb-cre* AAV injection on HFD for 10 weeks (n=5). Scale bar: 100 μm in **h** and **p**; 20 μm in the inset. Data are mean ± s.e.m. Statistical significance was calculated by unpaired Student’s two-sided t-test in all panels.

These findings were recapitulated in male LKO mice fed a Western diet (WD) (Fig. S2a-f) and, to a lesser extent, in female LKO mice on HFD, where the overall steatotic response was milder (Fig. S2g-k), consistent with the known sex differences in diet-induced hepatic lipid accumulation^43^.

To rule out developmental compensation in the constitutive LKO model, we implemented an inducible approach by injecting AAV8-albumin-cre to adult *Clstn3b^fl/fl^* mice and started a 10 weeks HFD feeding scheme 2 weeks after the injection. AAV-induced deletion of *Clstn3b* from hepatocytes exactly phenocopied the constitutive genetic KO model (Fig. 2i-p). These findings demonstrate that hepatocyte expression of CLSTN3B induced upon HFD feeding promotes lipid storage in the liver and the development of hepatic steatosis and injury, which can be alleviated by CLSTN3B ablation.

### Transgenic overexpression of CLSTN3B in hepatocytes promotes lipid accumulation and exacerbates hepatic steatosis

Our observation that ablation of hepatic CLSTN3B expression alleviates hepatic steatosis prompted us to examine whether overexpression of CLSTN3B is sufficient to promote lipid accumulation in hepatocytes. To this end, we generated a liver-specific CLSTN3B gain-of-function mouse model using a previously reported conditional *Clstn3b* transgenic allele (LTG: *albumin-cre*; *Clstn3b^tg/0^*)^38^. Even under the chow diet condition, the LTG mice exhibited significantly increased liver mass, liver-to-body weight ratio, and hepatic triglyceride content compared to the WT control (Fig. S3a-f). To further evaluate the impact of CLSTN3B overexpression on hepatic steatosis, we subjected the LTG mice and control littermates to HFD feeding. LTG mice showed a modest increase in liver size (p=0.09), significantly elevated liver-to-body weight ratio and hepatic triglyceride contents, and comparable body weight relative to controls (Fig. 3a-d). Although few visible LDs could be identified by histological analysis (Fig. 3e), the gross appearance of the liver and the thickened buoyant LD layer from liver homogenates both suggest increased hepatic lipid deposition in LTG mice (Fig. 3e).

**Figure 3.**
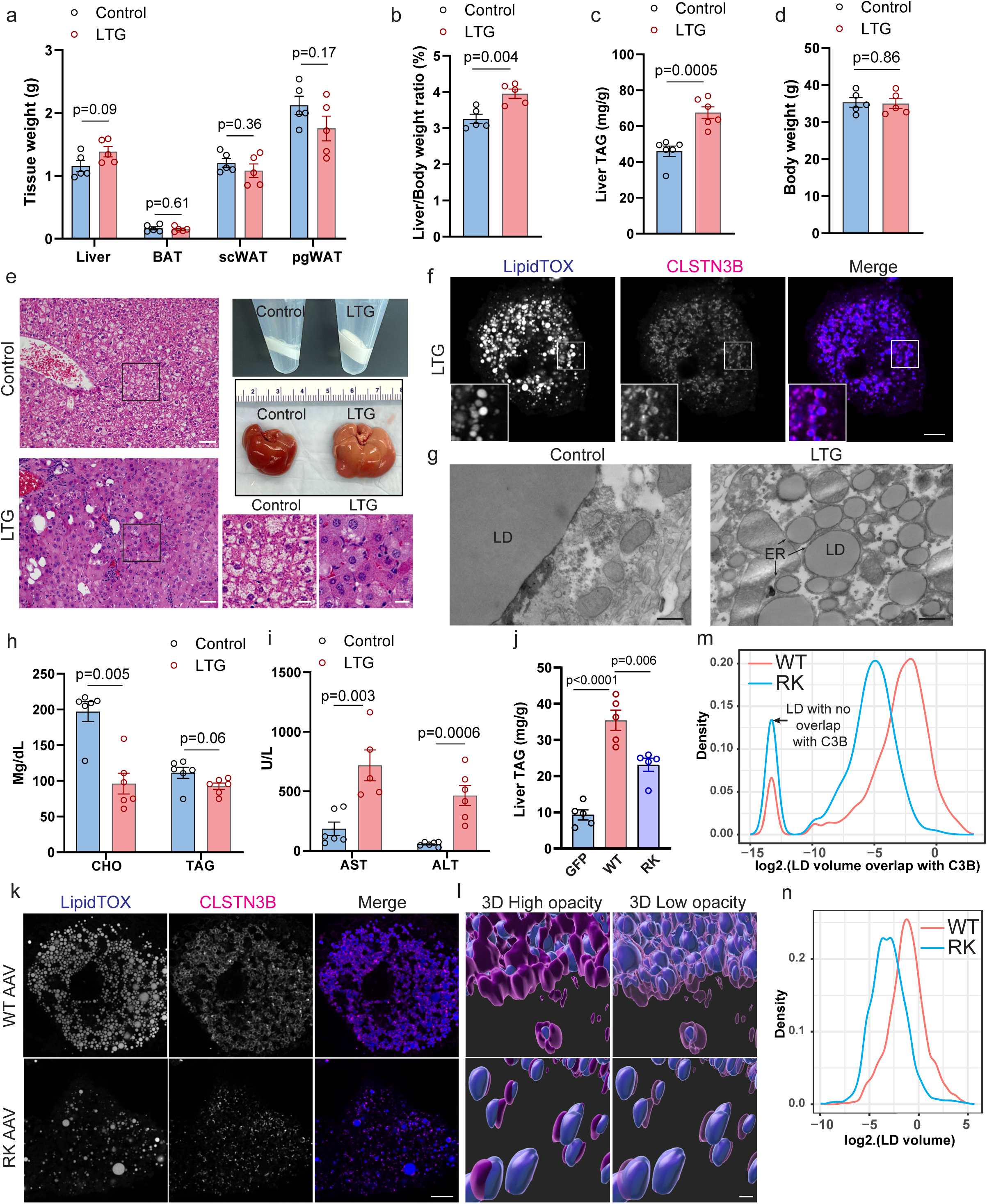
Transgenic overexpression of CLSTN3B in hepatocytes promotes lipid accumulation and exacerbates hepatic steatosis. **a**-**i**, Tissue weight (**a**), liver/body weight ratio (**b**), liver TAG content (**c**), body weight (**d**), liver gross appearance, buoyant LD fraction appearance and histology (**e**), fluorescence microscopy of primary hepatocytes (**f**), electron microscopy of liver sections (**g**), serum AST and ALT level (**h**), serum cholesterol and TAG level (**i**) from WT control and LTG mice on HFD for 10 weeks (n=5). **j**-**n**, Liver TAG content (**j**), fluorescence microscopy of primary hepatocytes (**k**), 3D reconstruction (**l**), LD/CLSTN3B contact volume distribution curve (**m**), and LD size distribution curve (**n**) in hepatocytes from mice receiving AAV-WT-CLSTN3B or AAV-RK-CLSTN3B injection. CLSTN3B surface opacity is adjusted in **l** to allow visualization of the LD surface enwrapped inside the CLSTN3B surface. The arrow in **m** denotes LDs not in contact with CLSTN3B signal but assigned with a small value to allow plotting on a logarithmic scale. Scale bar: 100 μm in **e**; 20 μm in the inset of **e**; 5 μm **f** and **k**; 500 nm in **g** and **l**. Data are mean ± s.e.m. Statistical significance was calculated by unpaired Student’s two-sided t-test in **a**, **b**, **c**,**d**, **h**, and **i,** and One-way ANOVA with Tukey’s post hoc test in **j**.

Fluorescence and electron microscopy revealed a large number of small LDs in LTG hepatocytes, characterized by robust CLSTN3B signals on the surface and extensive contact with the ER (Fig. 3f-g), mirroring previous observations made in adipocytes and other cell types^36^ and confirming those LDs are too small to be seen by standard H&E staining. Notably, serum triglyceride and cholesterol levels were reduced in the LTG mice (Fig. 3h), suggesting that CLSTN3B promotes intrahepatocellular lipid storage in cytosolic LDs at the expense of lipid output via the lipoprotein pathway. Substantial elevations in serum AST and ALT levels were observed, indicating exacerbated liver injury in the LTG mice (Fig. 3i). Similar results were observed with male LTG mice and control littermates on a WD (Fig. S3g-l) or Female LTG mice on a HFD (Fig. S3m-q).

To assess the molecular mechanism by which CLSTN3B promotes LD biogenesis and lipid storage in hepatocytes, we focused on the arginine-rich segment between its N-terminal LD-targeting and C-terminal ER-anchoring domains. We previously showed this segment facilitates hemifusion-like structures between the cytosolic leaflet of ER membrane and LD monolayer^36^. Replacement of 10 arginine residues with lysines (RK mutant) strongly impairs this activity. To test the functional importmance of the arginine-rich region *in vivo*, we used AAV to exogenously expresse WT CLSTN3B or the RK mutant in the livers of mice on chow diet. Despite comparable expression levels (Fig. S3s-t), mice receiving WT CLSTN3B accumulated significantly more hepatic triglycerides than those expressing the RK mutant (Fig. 3j). Fluorescence imaging with 3D reconstruction further revealed more extensive CLSTN3B localization on LD surfaces and larger LD sizes in WT-versus RK-expressing hepatocytes (Fig. 3k-n). Collectively, these data demonstrate that transgenic overexpression of CLSTN3B is sufficient to drive lipid storage and steatosis in hepatocytes, and that its arginine-rich segment is essential for promoting LD biogenesis and lipid accumulation *in vivo*.

### Hepatic CLSTN3B deficiency reduces LD surface phospholipid density and alters LD protein recruitment

The importance of the arginine-rich segment for CLSTN3B-driven LD biogenesis and lipid storage in hepatocytes suggests that CLSTN3B facilitates ER-to-LD phospholipid diffusion via hemifusion-like structures during LD expansion and maturation in hepatocytes, analogous to what we demonstrated in adipocytes. To directly assess this prediction, we quantified phosphatidylcholine (PC) and phosphatidylethanolamine (PE) densities on LD surfaces from primary hepatocytes following our previously published protocol^36^. CLSTN3B-deficient hepatocytes displayed significantly reduced LD surface PC and PE densities (Fig. 4a-b), consistent with impaired ER-to-LD phospholipid transfer. Reduced surface phospholipid coverage increases LD surface tension and promotes LD coalescence^44^. Accordingly, LD number and volume distributions in CLSTN3B-deficient hepatocytes were skewed toward larger droplets indicative of increased LD coalescence, as shown by both the LD number/size distribution and the weighted LD volume distribution curves (Fig. 4c-e).

**Figure 4.**
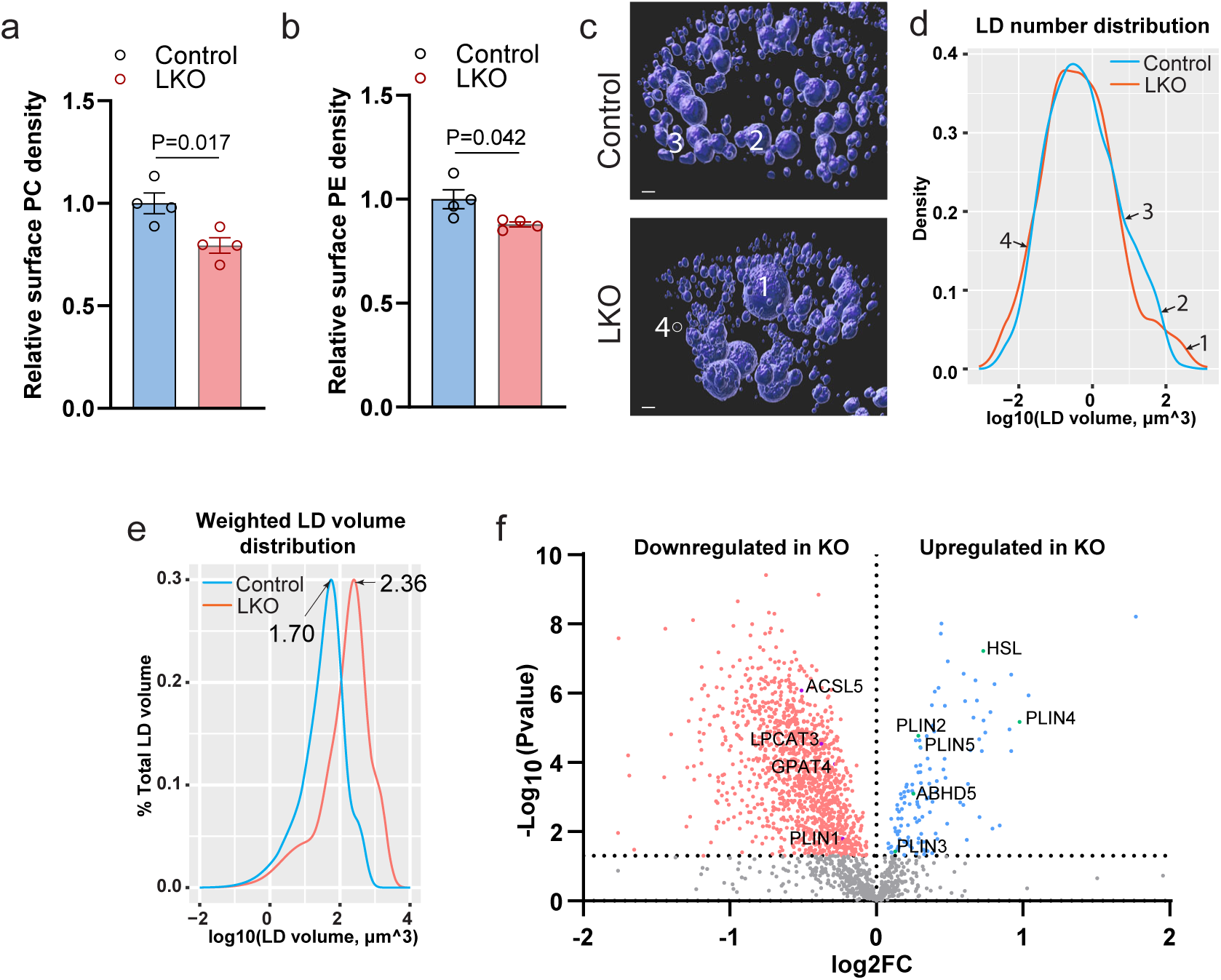
Hepatic CLSTN3B deficiency reduces LD surface phospholipid density and alters LD protein recruitment. **a**-**b**, Quantitation of surface PC (**a**) and PE (**b**) density on LDs isolated from WT and LKO mice liver (n=4). **c**-**e**, 3D reconstruction of fluorescence images (**c**), LD number/size distribution (**d**) and weighted LD volume distribution curve (**e**) of WT and LKO primary hepatocytes. The positions of 4 LDs of different sizes on the LD number/size distribution curve are labeled. **f**, Proteomics analysis of hepatocyte LDs isolated from WT and LKO mice (n=4). Scale bar: 3 μm in **c**. Data are mean ± s.e.m. Statistical significance was calculated by unpaired Student’s two-sided t-test in **a**, **b** and **f**.

Specifically, KO hepatocytes contained more LDs exceeding 100 μm^3^ (Fig. 4c-d, LD1), whereas WT hepatocytes showed a greater abundance of LDs in the 10–100 μm^3^ range (Fig. 4c-d, LD2 and 3). This pattern was recapitulated in the weighted LD volume distribution curves, which measures how triglyceride is partitioned among LDs of varying sizes (Fig. 4e). In WT hepatocytes, the curve peaked at a log_10_(LD volume) of 1.70 (∼50 μm^3^), whereas in KO hepatocytes, the peak shifted to 2.36 (∼230 μm^3^) (Fig. 4e). These findings with isolated primary hepatocytes also provided a quantitative characterization for the LD size difference observed between WT and LKO liver histology (Fig. 2h and p).

To understand how CLSTN3B impacts LD-associated protein recruitment in hepatocytes, we performed proteomics analysis of LDs isolated from the LKO or WT control hepatocytes.

All class II perilipins (PLIN2-5), which preferentially bind LDs with surface phospholipid packing defects, were found to be significantly upregulated in the LKO LD sample (Fig. 4f and Supplementary Table 1). In contrast, multiple class I proteins that access LDs via ER-LD membrane bridges, such as acyl-CoA synthetase long-chain family member 5 (ACSL5), lysophosphatidylcholine acyltransferase 3 (LPCAT3), and glycerol-3-phosphate acyltransferase 4 (GPAT4)^14^, were significantly downregulated on the LKO LDs (Fig. 4f), Distinct from all class II perilipins, PLIN1, which was recently shown to behave as a class I protein^45^, was found to be downregulated on the LKO LD (Fig. 4f), an observation that we also made in *Clstn3b^-/-^* adipocytes^36^. Together, the increased abundance of class II proteins and reduced abundance of class I proteins further support the notion that CLSTN3B promotes the hemifusion-like structure between the ER and LD.

### Hepatic CLSTN3B deficiency increases energy expenditure and reduces metabolic efficiency

Our finding that the LKO mice accumulate less lipid in the liver without concomitantly increased lipid export prompted us to investigate whether loss of CLSTN3B alters systemic energy balance. We thus performed comprehensive metabolic profiling of LKO and WT control littermates. Despite comparable food intake (Fig. 5a), LKO mice exhibited significantly slower weight gain upon HFD feeding (Fig. 5b, Two-way ANOVA, p_genotype x time_=0.018) without significant difference in body composition (Fig. S4a). This phenotype was recapitulated with the AAV-induced liver-specific CLSTN3B KO cohort (Fig. 5c, Two-way ANOVA, p_genotype x time_=0.0069), suggesting a hepatocyte-intrinsic role for CLSTN3B in regulating systemic energy metabolism. Indirect calorimetry revealed significantly higher O_2_ consumption, CO_2_ production, and heat production in LKO mice on HFD for 8 weeks than the control mice (Fig. 5d-f), without notable differences in locomotive activity (Fig. S4b). Rectal temperature was also significantly higher in LKO mice at ZT18 (Fig. 5g). Together, these findings indicate that the LKO mice have increased energy expenditure and reduced metabolic efficiency.

**Figure 5.**
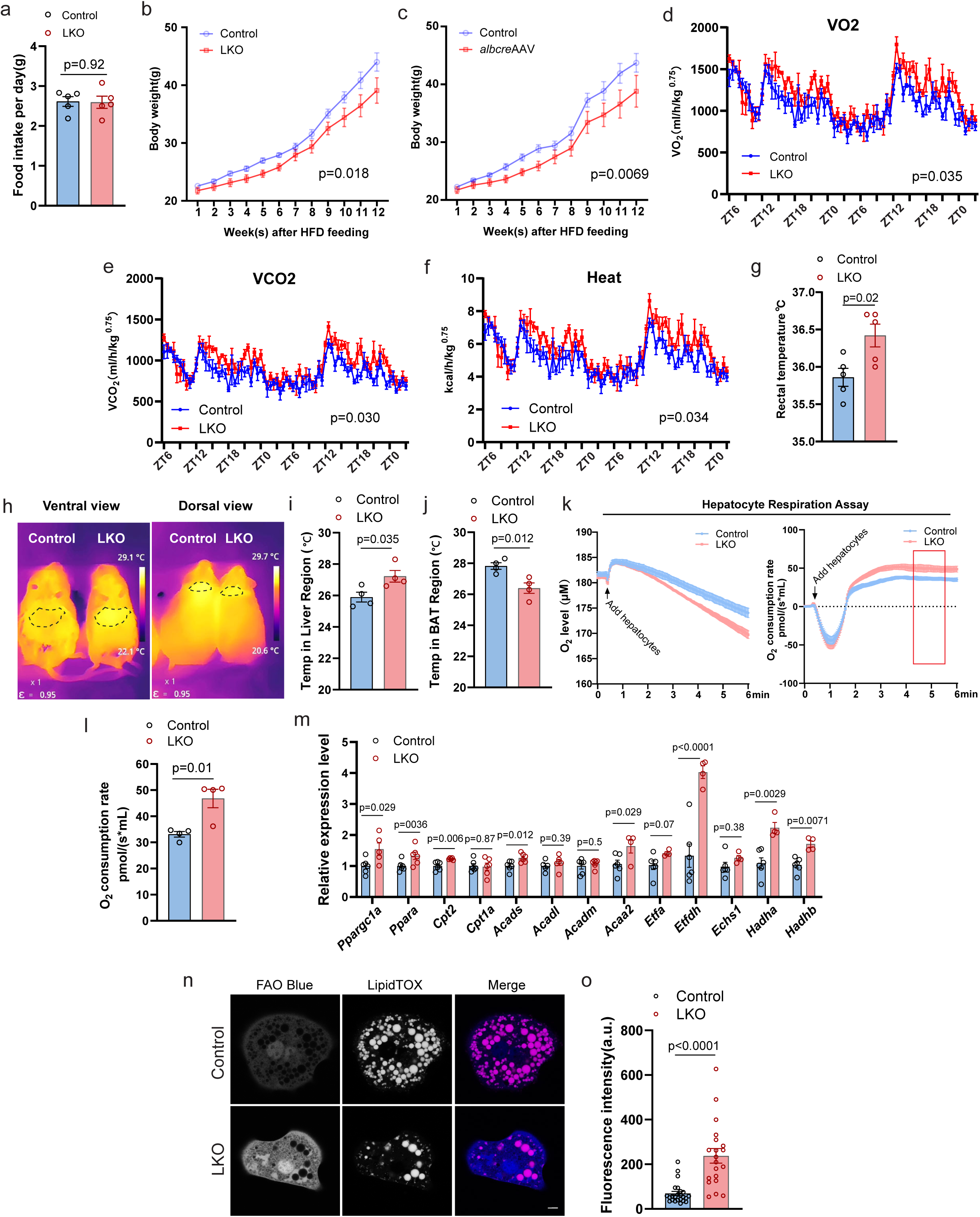
Hepatic CLSTN3B deficiency increases energy expenditure and decreases metabolic efficiency. **a**, Daily food intake of WT control and LKO mice (n=5). **b**-**c**, Body weight growth curve of a WT control and LKO cohort (n=5) (**b**) or an AAV-induced liver-specific CLSTN3B KO cohort (n=5) (**c**) maintained on HFD. **d**-**f**, Indirect calorimetry analysis of WT control and LKO mice on HFD for 10 weeks (n=5). O_2_ consumption rate (**d**), CO_2_ production rate (**e**), and calculated heat production rate (**f**) normalized to body weight raised to the 0.75 power over a 45-hour period are shown. **g**, Rectal temperature of WT and LKO mice at ZT18 (n=5). **h**-**j**, Infrared thermography images (**h**) and quantitation of surface temperature in the liver (**i**) and BAT region (**j**) of WT and LKO mice (n=3-4). **k**-**l**, Respiration assay of freshly isolated primary hepatocytes from WT and LKO mice. O_2_ concentration and the time derivative traces (**k**) and quantitation of O_2_ consumption rate are shown (**l**) (n=4). The bracket in **k** denotes the time window during which O_2_ consumption rate was sampled. **m**, qPCR analysis of hepatic mitochondrial biogenesis and FAO gene expression in WT and LKO mice (n=5). **n**-**o**, Fluorescence image (**n**) and quantitation (**o**) of FAO blue assay on primary WT and LKO hepatocytes (n=20-24). Data are mean ± s.e.m. Statistical significance was calculated by Two-way Repeated Measurement ANOVA in **b**, **c**, **d**, **e**, and **f**, and unpaired Student’s two-sided t-test in **g**, **i**, **j**, **l**, **m**, and **o**.

To identify the primary organ responsible for the increased energy expenditure, we used infrared thermography. LKO mice showed higher surface temperature over the liver region but not over brown adipose tissue (BAT), which instead trended cooler (Fig. 5h-j), suggesting increased hepatic metabolic activity. In support of this, primary hepatocytes isolated from LKO mice exhibited significantly elevated respiration rates compared to control cells (Fig. 5k-l), consistent with increased liver thermogenesis. The LTG mice showed exactly the opposite phenotypes, including lower surface temperature over the liver region and reduced primary hepatocyte oxygen consumption rate (Fig. S4c-f)

Given the combination of elevated respiration and reduced lipid accumulation in LKO hepatocytes, we hypothesized that hepatic FAO is enhanced. Indeed, gene expression analysis revealed that the mRNA levels of peroxisome proliferator-activated receptor gamma coactivator 1-alpha (PGC1α), a key promoter of mitochondrial biogenesis, and PPARα, the nuclear receptor regulating genes involved in FAO, along with multiple PPARα target genes involved in FAO are upregulated in the LKO liver (Fig. 5m). FAO Blue fluorescence assays confirmed increased FAO enzymatic activity in LKO hepatocytes (Fig. 5n-o).

Interestingly, reduced surface temperature in the BAT region, along with increased lipid deposition and downregulation of thermogenic genes in the BAT of LKO mice (Fig. S4g-h), suggests suppressed BAT thermogenesis. This may reflect compensatory downregulation of sympathetic tone to the BAT due to excess hepatic heat production.

Together, these findings across organismal, organ, and cellular levels demonstrate that hepatic CLSTN3B deficiency promotes liver FAO, elevates whole-body energy expenditure, and reduces metabolic efficiency.

### Hepatic CLSTN3B deficiency drives a lipolysis/re-esterification futile cycle

We next sought to understand the mechanism underlying elevated FAO and O_2_ consumption displayed by CLSTN3B-deficient hepatocytes. Mitochondrial uncoupling or AMP-activated protein kinase (AMPK) activation are known to enhance FAO. Nevertheless, CLSTN3B-deficient hepatocytes exhibited *increased* mitochondrial membrane potential, as detected by tetramethylrhodamine ethyl ester (TMRE) staining, and a *higher* ATP/ADP ratio compared to control hepatocytes (Fig. 6a-c), indicating more active coupled respiration/oxidative phosphorylation and no energy deficiency. Contraty to the KO cells, CLSTN3B-transgenic hepatocytes displayed significantly decreased mitochondrial membrane potential, consistent with their reduced respiration rate (Fig. S5a-b). These findings thus argue against mitochondrial uncoupling or AMPK activation as primary mechanisms driving increased FAO of CLSTN3B-deficient hepatocytes.

**Figure 6.**
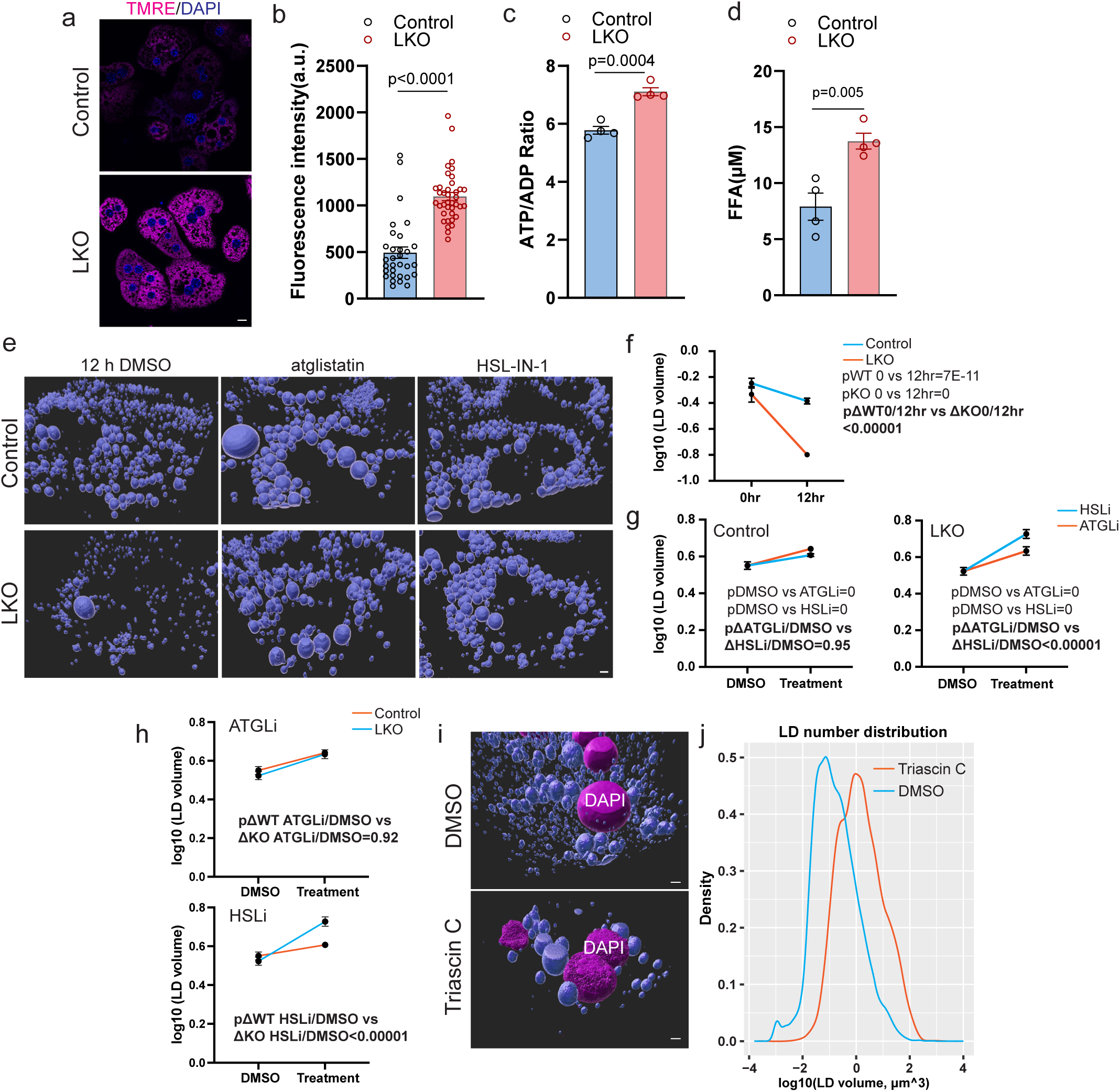
Hepatic CLSTN3B promotes a lipolysis/re-esterification futile cycle. **a**-**b**, Fluorescence image (**a**) and quantitation (**b**) of mitochondrial membrane potential measured by TMRE staining in WT and LKO primary hepatocytes (n=31-38). **c**, ATP/ADP ratio measurement in WT and LKO primary hepatocytes (n=4 biological replicates). **d**, Lipolysis assay of WT and LKO primary hepatocytes (n=4 biological replicates). **e**-**h**, 3D reconstruction of fluorescence images (**e**), comparison of LD sizes in WT and LKO hepatocytes between 0 hr or 12 hr (**f**) or treated with different lipase inhibitors (**g**) and (**h**). **i**-**j**, 3D reconstruction of fluorescence images (**i**), LD number/size distribution curve (**j**) of LKO primary hepatocytes treated with DMSO or triascin C. Scale bar: 5 μm in **a**; 3 μm in **e** and **i**. Data are mean ± s.e.m. Statistical significance was calculated by unpaired Student’s two-sided t-test in **b**, **c**, and **d**, and One-way ANOVA with Tukey’s correction for p values in regular type in **f** and **g**, and One-way ANOVA with Tukey’s correction based on the sample size, means, and standard errors derived from Two-way ANOVA with Tukey’s correction for p values in bold in **f**, **g**, and **h**.

Aside from mitochondrial uncoupling, futile metabolic cycles, such as what was described previously in the BAT or cancer cells^46–49^, can also increase energy expenditure. We became interested in the lipolysis/re-esterification cycle because we previously observed increased lipolysis in *Clstn3b^-/-^* adipocytes^36^. To examine whether CLSTN3B deficiency leads to increased lipolysis in hepatocytes, we treated primary hepatocytes with etomoxir and triascin C to block FFA oxidation and re-esterification and measured FFA released into the extracellular space. The LKO hepatocytes released significantly more FFAs than control cells, whereas the LTG cells showed the opposite (Fig. 6d, Fig. S5c), confirming that CLSTN3B deficiency promotes hepatocyte lipolysis.

Interestingly, the ATGL activator ABHD5, and the lipase hormone sensitive lipiase (HSL), are significantly upregulated on the LKO LD (Fig. 4f). These proteins recognize packing defects through bulky hydrophobic residues^50,51^, and their recruitment likely contributes to the enhanced lipolysis observed in CLSTN3B-deficient hepatocytes. To evaluate their functional relevance, we treated WT and CLSTN3B-deficient hepatocytes with inhibitors of ATGL (Atglistatin) or HSL (HSL-IN-1) *in vitro*. Over a 12-hour period, LD sizes in vehicle treated cells decreased markedly (Fig. 6e-f. See supplementary table 2 for complete statistical analysis), with the magnitude of reduction being more pronounced in CLSTN3B-deficient hepatocytes than in control cells (Fig. 6g, p value in bold reflects the significance of the difference between pair-wise comparisons), confirming more extensive lipolysis in the KO cells. Both ATGL or HSL inhibition significantly increased LD size relative to the vehicle control in WT and LKO hepatocytes (Fig. 6e and g). Notably, HSL inhibition was significantly more effective than ATGL inhibition at enlarging LD sizes in LKO but not WT hepatocytes (Fig. 6g, p value in bold). The magnitude of LD enlargement induced by HSL inhibition is also significantly greater in LKO than in WT hepatocytes (Fig. 6h, p value in bold). Similar trends could be visualized in the weighted LD size distribution curves (Fig. S5d-e). These findings are consistent with the increased recruitment of HSL to LDs in the absence of CLSTN3B and support a pronounced functional importance of HSL in driving lipolysis in LKO cells.

FFA re-esterification is required to complete the futile cycle. Curiously, aside from a few large LDs, CLSTN3B-deficient hepatocytes also contain numerous small LDs (<0.1 μm^3^) (Fig. 4c-d, LD4), which may be newly formed from re-esterified FFA released from lipolysis, similar to what occurs in adipocytes under norepinephrine (NE) stimulation^36,52^. To directly test this possibility, we treated CLSTN3B-deficient hepatocytes with triascin C, a small molecule inhibitor of ACSL enzymes catalyzing fatty-acyl CoA formation, the first step in re-esterification, over a 12-hour period. This treatment led to a marked depletion of the small LD population (<0.1 μm^3^) (Fig. 6i-j), confirming that these small LDs arise from re-esterification of lipolysis-derived FFAs.

Together, these findings reveal that CLSTN3B deficiency in hepatocytes promotes a futile lipolysis/re-esterification cycle, explaining chronically elevated FAO and energy expenditure and reduced metabolic efficiency displayed by the LKO mice.

### Hepatic CLSTN3B Deficiency Attenuates Steatosis-Induced Fibrosis

Hepatic steatosis is a known driver of hepatocyte injury, inflammation, HSC activation, and ultimately fibrosis, a central pathological feature of MASLD. Given our findings that hepatocyte-specific deletion of CLSTN3B ameliorates hepatic steatosis in HFD-fed mice, we hypothesized that CLSTN3B ablation might also confer protection against hepatic fibrosis. To test this, we subjected the LKO and LTG mice, along with their respective control mice, to a WD for 16 weeks, a regimen that more robustly models MASLD pathology than HFD alone. Fluorescence imaging revealed markedly reduced α-SMA and F4/80 staining in LKO liver sections compared to controls (Fig. 7a-d), indicating attenuated hepatic fibrosis and decreased macrophage infiltration. Consistently, Sirius Red staining confirmed lower collagen deposition in LKO livers (Fig. 7e). In contrast, LTG mice displayed exacerbated fibrosis and increased macrophage infiltration relative to controls (Fig. 7a-d). Gene expression analyses further supported these histological findings: LKO livers exhibited significantly reduced expression of pro-fibrotic and pro-inflammatory genes, while LTG livers showed elevated levels of these transcripts (Fig. 7f-g).

**Figure 7.**
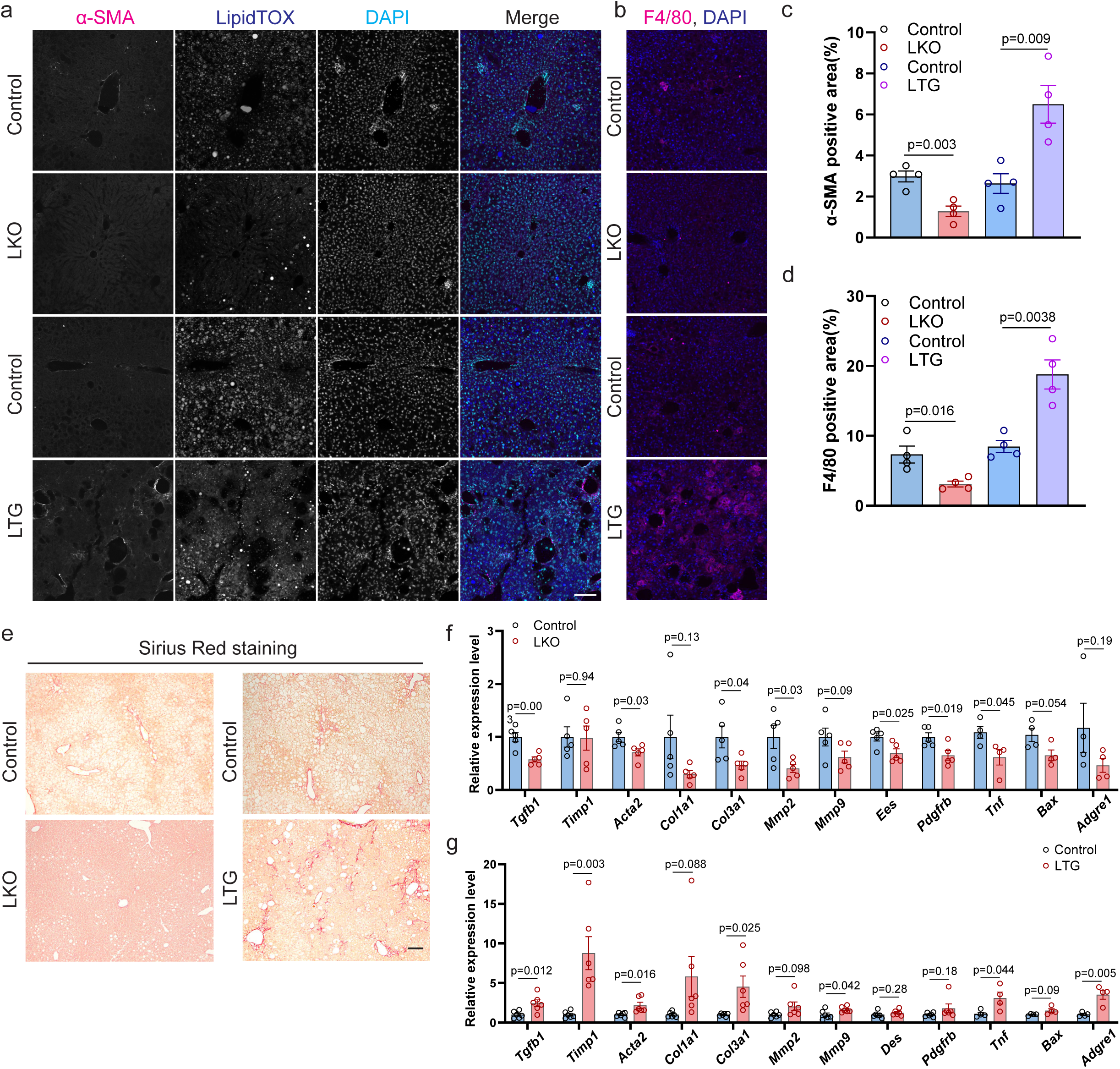
Hepatic CLSTN3B deficiency attenuates steatosis-induced fibrosis. **a**-**d**, Immunostaining of the fibrosis marker α-SMA and the macrophage marker F4/80 in liver sections of control, LKO, and LTG mice (n=4 biological replicates). Fluorescence images of α-SMA (**a**) and F4/80 (**b**), and quantitation of α-SMA (**c**) and F4/80 (**d**) are shown. **e**, Sirium Red staining of liver sections of control, LKO, and LTG mice. **f**-**g**, qPCR analysis of genes associated with hepatic fibrosis and inflammation in the liver of WT control and LKO mice (**f**) and WT control and LTG mice (**g**) (n=4-5). Scale bar: 100 μm in **a**, **b**, and **e**. Data are mean ± s.e.m. Statistical significance was calculated by unpaired Student’s two-sided t-test in all panels.

To determine whether these phenotypes reflect CLSTN3B function during dietary invention and disease progression rather than developmental consequences, we acutely modulated hepatic CLSTN3B levels in WD-fed WT mice using AAV-mediated gene delivery. Indeed, AAV-induced CLSTN3B knockdown or overexpression phenocopied the respective LKO and LTG models (Fig. S6a-d), confirming that CLSTN3B activity during WD feeding is critical for regulating steatohepatitis and fibrosis.

Taken together, these results demonstrate that hepatocyte-specific deletion of CLSTN3B significantly mitigates the progression of steatohepatitis and fibrosis, whereas transgenic overexpression of CLSTN3B aggravates these pathological features.

### Hepatic *CLSTN3B* expression level positively correlates with clinical MASLD fibrosis severity and progression

We next sought to determine clinical relevance of hepatic *CLSTN3B* expression in MASLD, which is characterized by progressive liver fibrosis due to chronic steatohepatitis, in clinical cohorts of MASLD patients. First, we developed a method to accurately quantitate *CLSTN3B* expression without interference from the *CLSTN3* gene. This is particularly important for analyzing human data because *CLSTN3* is expressed at a substantially higher level than *CLSTN3B* in human liver. Per the GENCODE annotation release 45, *CLSTN3B* is defined as one of the *CLSTN3* transcripts (ENST00000535313.2). *CLSTN3B* comprises three exons, with the first exon being unique. The second and third exons are shared with *CLSTN3*. To differentiate *CLSTN3B* expression from *CLSTN3*, we removed two shared exons (chr12: 7,157,489-7,157,691, chr12: 7,157,941-7,158,305) from either gene records. This left only the first exon of *CLSTN3B* (chr12: 7,155,868-7,156,929) and the remaining exons of *CLSTN3* to allow us to separately and accurately quantitate *CLSTN3B* and *CLSTN3* expression.

With this method, we found that both *CLSTN3B* and *CLSTN3* expression were higher in 206 European MASLD patients compared to 10 healthy obese controls (p=0.000068 and p=0.0017, respectively) (Fig. 8a, Fig. S7a). Of note, expression levels of *CLSTN3B*, not *CLSTN3*, were positively correlated with higher liver fibrosis stage (Spearman’s rho=0.16, p=0.023) (Fig. 8b, Fig. S7b). Similar correlation of *CLSTN3B* with fibrosis stage was observed in 106 non-cirrhotic Japanese MASLD patients (p=0.03) (Supplementary Table 3). Among them, 77 patients had paired liver biopsy specimens over the course of longitudinal clinical follow-up (median interval, 2.3 years). Time-adjusted progression of liver fibrosis stage was positively correlated with elevated baseline *CLSTN3B*, not *CLSTN3*, expression levels (Spearman’s rho=0.26, p=0.023) (Fig. 8c, Fig. S7c). Notably, in this Japanese cohort, a non-significant trend (p=0.055, Fisher test) was observed in the correlation between steatosis severity and *CLSTN3B* expression level when the comparison was made between the bottom quartile (Q1) of *CLSTN3B* expression level and the rest (Q2-Q4), suggesting a potential impact of *CLSTN3B* expression on steatosis in MASLD patients. The hepatic *CLSTN3B* expression was positively correlated with *CLSTN3* expression (Fig. S7d, Spearman’s rho=0.29, p=0.023), but the association with fibrosis progression is unique to *CLSTN3B* and not affected by *CLSTN3* (Supplementary Table 4).

**Figure 8.**
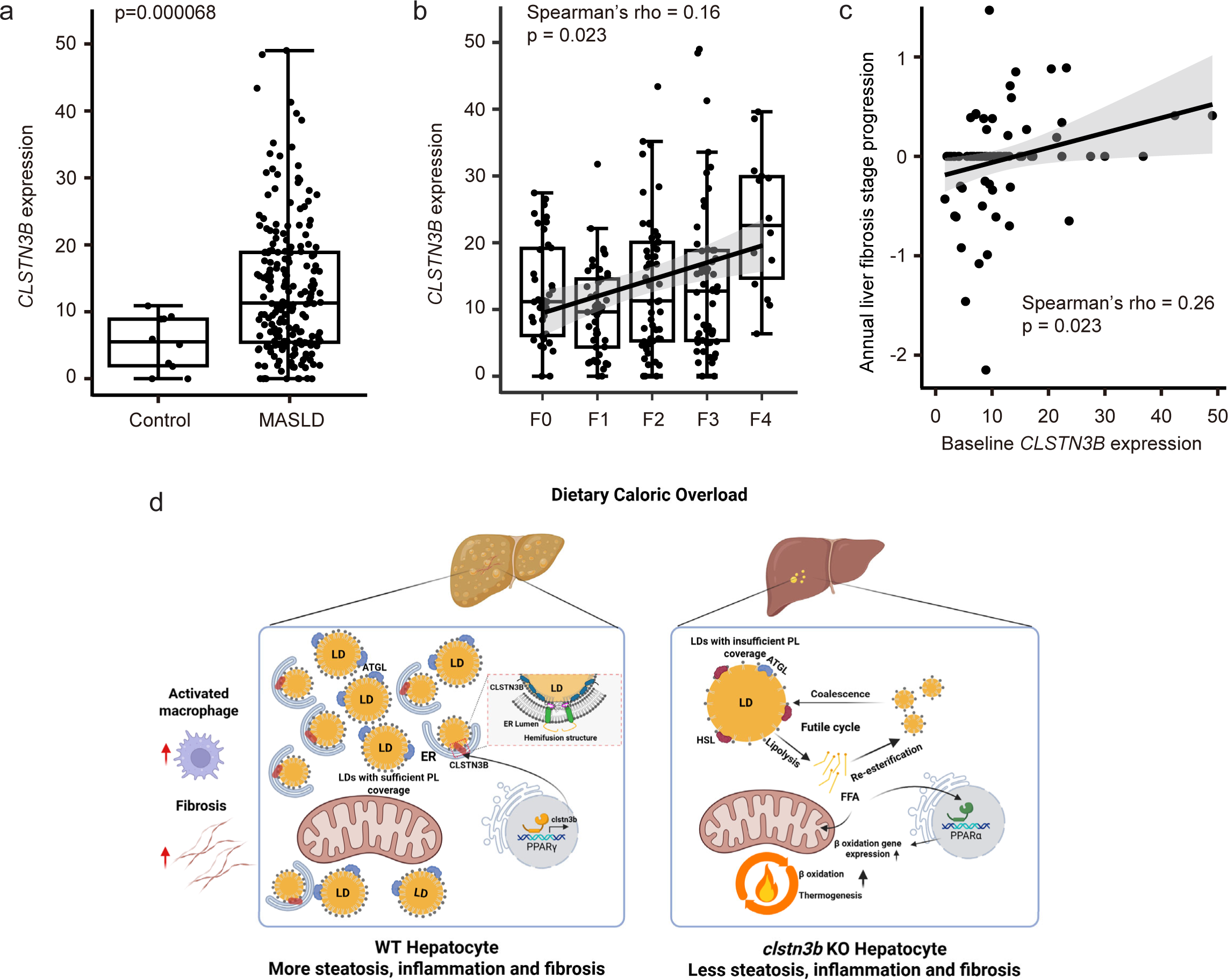
Hepatic *CLSTN3B* expression level positively correlates with clinical MASLD progression. **a**, Hepatic *CLSTN3B* expression levels in 206 European MASLD patients compared to 10 healthy obese controls. **b**, Hepatic *CLSTN3B* expression levels by liver fibrosis (F) stage in 206 European MASLD patients. **c**, Correlation between baseline *CLSTN3B* expression levels and time-adjusted (i.e., annual) progression of liver fibrosis stage in 77 Japanese MASLD patients. Inter-group difference was assessed by Wilcoxon rank-sum test. Correlation was assessed by Spearman correlation test. **d**, Model of CLSTN3B’s effect on LD dynamics in hepatocytes.

## Discussion

Our study identifies CLSTN3B as a previously unrecognized regulator of LD biogenesis and systemic energy homeostasis in hepatocytes, whose expression is strongly induced by dietary caloric overload via PPARγ activation. Through both gain- and loss-of-function mouse models, we demonstrate that CLSTN3B facilitates hepatic lipid storage by promoting phospholipid transfer from the ER to expanding LDs and also supporting ER-to-LD protein translocation. In the absence of CLSTN3B, hepatocyte LDs exhibit reduced phospholipid surface coverage and enhanced recruitment of cytosolic LD-targeting proteins, including HSL, which drives a robust lipolysis/re-esterification futile cycle (summarized in Fig. 8d). This metabolic shift increases FAO, reduces hepatic triglyceride accumulation, and ultimately enhances whole-body energy expenditure while conferring resistance to MASLD-associated pathology. Consistent with these findings, clinical data reveal a positive correlation between hepatic CLSTN3B expression and both liver fibrosis severity and progression in MASLD patients. Collectively, our results position CLSTN3B as a critical molecular link between nutrient excess and hepatocellular lipid handling, with broad implications for metabolic efficiency, energy balance, and susceptibility to MASLD.

Our findings in hepatocytes reinforce prior conclusions from the adipocyte system, underscoring the core function of CLSTN3B as a key regulator of lipid droplet (LD) maturation by facilitating phospholipid and protein trafficking via the hemifusion-like structure between ER and LD stabilized by its arginine-rich segment. This may seem paradoxical, as CLSTN3B is most abundantly expressed in brown adipocytes, which are better known for FAO and thermogenesis rather than lipid storage. However, LD maturation is critical for shaping the lipolytic response of brown adipocytes: immature LDs exhibit elevated basal lipolysis but reduced responsiveness to hormonal stimulation^36^. Moreover, CLSTN3B promotes sympathetic innervation of brown adipocytes via an S100B-dependent mechanism^38^, a function not observed in white adipocytes or hepatocytes, which receive minimal direct sympathetic input. Therefore, in white adipocytes and hepatocytes, where neural stimulation of lipid mobilization is limited, the primary manifestation of CLSTN3B-mediated LD maturation appears to be enhancement of lipid storage capacity. Interestingly, a recent study reported that CLSTN3B is upregulated in mouse hepatocytes in a PPARα-dependent manner during ketogenic diet (KD) feeding and is required for efficient FAO and ketogenesis^42^. This observation seems at odds with our finding that CLSTN3B suppresses FAO under high-fat diet (HFD) or Western diet (WD) conditions. However, it is plausible that KD creates a distinct metabolic environment in hepatocytes, representing another case of context-dependent outcome of LD maturation. For example, structurally mature LDs may facilitate more efficient fatty acid channeling to mitochondria under KD conditions, but not under HFD or WD, where mitochondrial function and lipid trafficking may be compromised. To resolve this apparent discrepancy, a more detailed comparison of LD composition and inter-organelle communication across dietary contexts will be necessary.

Our findings align with previous studies showing that deficiencies in LD-associated proteins such as PLIN2 and CIDEB impair LD biogenesis and lipid storage, thereby mitigating the onset of MASLD pathologies^27–32^. Notably, while CLSTN3B-deficient hepatocytes display fewer but conspicuously large LDs, the loss of PLIN2 or CIDEB results in a more marked reduction in LD size, suggesting a more severe disruption of LD biogenesis in those contexts. It remains to be seen whether the steatosis-alleviating effects observed in PLIN2 or CIDEB deficiency are mediated by an enhanced lipolysis/re-esterification cycle, as we demonstrate for CLSTN3B deficiency. However, at least one study has reported increased energy expenditure following PLIN2 knockdown^32^, hinting at a potentially shared mechanism.

The increased recruitment of HSL and its functional significance in CLSTN3B-deficient hepatocytes is also noteworthy. Under normal physiological conditions, ATGL is considered the primary driver of hepatic lipolysis^53^, whereas HSL, by contrast, has been thought to primarily hydrolyze cholesterol esters rather than triglycerides in hepatocytes^54^. However, HSL expression is induced by PPARγ in hepatocytes^55^, and may thus gain functional significance in the liver upon HFD feeding. Indeed, HSL inhibition yielded a comparable effect to ATGL inhibition at enlarging LDs in WT hepatocytes, suggesting that HSL, when expressed at a higher level, may become more relevant as a triglyceride lipase. Furthermore, our findings suggest that in the presence of structurally aberrant LDs, such as those formed in the absence of CLSTN3B, HSL may gain additionally enhanced access to the triglyceride core and contribute more significantly to triglyceride catabolism. Supporting this possibility, previous work has shown that adenovirus-mediated overexpression of HSL in the liver markedly reduces hepatic triglyceride accumulation^56^. This indicates that HSL, although not typically considered a major hepatic triglyceride hydrolase under physiological conditions, could acquire therapeutic relevance in specific contexts. Further investigation is warranted to determine whether a similar shift in HSL activity occurs in PLIN2- or CIDEB-deficient settings and whether this contributes to their respective anti-steatotic phenotypes.

Among the five perilipins, PLIN1 is unique in exhibiting reduced abundance on LDs from CLSTN3B-deficient hepatocytes. This suggests that PLIN1 targeting to LDs may involve a mechanism distinct from the phospholipid packing-sensing mode used by other perilipins via amphipathic helices. Two possible mechanisms may underlie this observation. First, rather than competing with phospholipids for LD surface space, PLIN1 may bind cooperatively with them. Supporting this idea, previous studies have shown that (1) PLIN1 retention on artificial LDs is enhanced by co-packing with phospholipids^57^, and (2) PLIN1, but not PLIN2 or PLIN3, can associate with artificial LDs fully coated with phospholipids^58^. This cooperative interaction may reflect PLIN1’s greater hydrophobicity, deeper insertion into the phospholipid monolayer, and potential engagement with phospholipid acyl chains. In the absence of CLSTN3B, reduced phospholipid coverage on LDs may disrupt these interactions, diminishing PLIN1 retention as observed in our FRAP assays using *Clstn3b-/-* adipocytes^36^. Second, PLIN1 has been shown to behave like a class I LD protein, embedding in the ER membrane in the absence of LDs, and may thus require ER-LD membrane bridges for translocation^45^. Since CLSTN3B facilitates the formation and stabilization of hemifusion-like ER-LD connections, its loss likely impedes PLIN1 translocation from the ER to LDs, and cause the reduced abundance of PLIN1 on LDs.

The increased TMRE fluorescence intensity and elevated ATP/ADP ratio observed in CLSTN3B-deficient hepatocytes point to an intriguing link between CLSTN3B-mediated LD maturation and mitochondrial function. These findings suggest that alterations in LD surface composition, driven by CLSTN3B ablation, may impact LD-mitochondria interactions and influence mitochondrial activity under the HFD or WD-induced steatosis condition. Notably, PLIN5, a protein known to localize at LD-mitochondria contact sites^59^, is significantly upregulated on LDs in the absence of CLSTN3B, implying that LD-mitochondria coupling may be enhanced under these conditions. Two distinct mitochondrial subpopulations with specialized functions have been described: cytosolic mitochondria (CM), which are more active in FAO, and peridroplet mitochondria (PDM), which preferentially support lipid synthesis^60^. Prior studies have reported divergent roles for PLIN5, with some suggesting it promotes lipid storage, while others propose it facilitates fatty acid transfer from LDs to mitochondria for β-oxidation^61–64^. Both roles could plausibly coexist in CLSTN3B-deficient hepatocytes. For example, PLIN5 may facilitate FFA transfer from large, lipolytically active LDs to CM for FAO, while simultaneously supporting PDM activity near small, nascent LDs by promoting re-esterification and maintaining futile cycling. To discern between these non-mutually exclusive functions, it will be important to determine whether PLIN5 preferentially associates with large versus small LD populations. Such spatial analysis could provide critical insights into the dominant function of PLIN5 in the context of CLSTN3B deficiency.

Further complicating the picture, recent evidence suggests that post-translational modifications of PLIN5 may govern its functional switch between lipid storage and lipolysis^65^. Whether CLSTN3B influences PLIN5 through such modifications remains an open and compelling question for future investigation.

Our finding that CLSTN3B overexpression promotes LD biogenesis and triglyceride storage while simultaneously exacerbating MASLD-related pathologies, including steatosis, liver injury, and fibrosis, offers new insights into the mechanisms linking hepatocyte lipid metabolism and MASLD progression. Although lipotoxicity is often implicated in MASLD, our previous findings and the current study all suggest that CLSTN3B promotes extensive association between ER and LD, which strongly excludes LD-targeted protein and suppresses lipolysis^36^. This condition would conceivably prevent the accumulation of toxic lipid intermediates, which are commonly derived from partial triglyceride hydrolysis or unesterified FFAs. Therefore, alternative mechanisms beyond classical lipotoxicity may underlie the worsening of MASLD phenotypes in the context of CLSTN3B overexpression. Recent studies have implicated mitochondrial dysfunction and cardiolipin deficiency in MASLD pathogenesis^66^. Cardiolipin synthesis in mitochondria relies on the transfer of phosphatidic acid (PA) from the ER through ER-mitochondria contact sites^67^. We propose that excessive LD biogenesis, particularly under conditions of CLSTN3B overexpression, may disrupt this lipid trafficking pathway by diverting PA-rich ER membranes toward LD formation at the expense of mitochondrial PA delivery. Notably, we showed that CLSTN3B preferentially utilizes PA to promote hemifusion-like ER-LD membrane bridges^36^, further raising the possibility that CLSTN3B overexpression reduces PA availability for cardiolipin biosynthesis in mitochondria. This phospholipid redistribution may impair mitochondrial electron transport chain (ETC) function and promote reactive oxygen species (ROS) production, thereby contributing to MASLD pathology through mitochondrial stress rather than lipotoxicity. Future studies are warranted to determine whether enhanced LD biogenesis directly limits mitochondrial PA supply, disrupts cardiolipin synthesis, and impairs ETC function, ultimately promoting oxidative stress and liver injury in MASLD.

## Methods

### Mouse strains

All animal studies were approved by and in full compliance with the ethical regulation of the Institutional Animal Care and Use Committee (IACUC) of University of Texas Southwestern Medical Center. Mice were euthanized by CO_2_ asphyxiation, according to the guidelines of IACUC and the recommendations of the Panel on Euthanasia of the American Veterinary Association.

The *Clstn3b flox/flox* and the *Clstn3b* transgene mice were described in a previous publication (Zhang et al, PMID: 38293096). Mice were maintained on a normal chow diet (13% fat, 57% carbohydrate and 30% protein, PicoLab Rodent Diet 5L0D) or placed on a high-fat diet (HFD, Rodent diet with 60 kcal% Fat, Research diet, D12492, USA) or western diet consisted of high sugar, high fat, and high cholesterol solid food (Teklad Diets #TD. 120528) combined with high-sugar water containing 23.1 g/L d-fructose (Sigma-Aldrich #G8270) and 18.9 g/L d-glucose (Sigma-Aldrich #F0127) or CDA-HFD consisted of a methionine and choline-deficient high-fat diet (Research Diets #A06071302) from 6-8 weeks of age for 8 to 16 weeks. All mice were maintained under a 12 hr light/12 hr dark cycle at constant temperature (23°C or 30°C as specified) and 40–60% humidity with free access to food and water. Liver specific Clstn3b knockout mice were generated via crossing *Clstn3b flox/flox* mice with *Albcre* mice. Liver specific Clstn3b transgenic mice were generated via crossing *Clstn3b* transgene mice with *Albcre* mice. All animal studies were approved by and in full compliance with the ethical regulation of the Institutional Animal Care and Use Committee of University of Texas Southwestern Medical Center. Sample size was chosen based on literature and pilot experiment results to ensure that statistical significance could be reached. Randomization was not performed because mice were grouped based on genotype. Littermates were used for all the experiments involving the conditional KO and transgenic mice.

### AAV production and purification

AAV production and purification were described in a previous publication (Wang et al, PMID: 37040760). Briefly, AAV8 was produced using AAV-Pro 293T cells (Takara #632273) cultured in one or more 15cm dishes. Cells were plated one day before transfection at 50% confluence, which would allow the cells to reach 80-90% confluence the next day. For transfection of one 15cm dish, 10mg MOSAICS vector, 10mg pAAV2/8 (Addgene #112864), and 20mg pAdDeltaF6 (Addgene #112867) plasmids were mixed with 1ml Opti-MEM medium in one tube. In another tube, 160ml PEI solution (1mg/ml in water, pH7.0, powder from ChemCruz #sc-360988) was mixed with 1ml Opti-MEM medium. The solutions from both tubes were then mixed and incubated for 10min before adding to cell culture. 48h after transfection, the cells were scraped off the dish and collected by centrifugation at 500g for 10min. The supernatant was disinfected and discarded, and the cell pellets were lysed in 1.5ml/15cm dish lysis buffer (PBS supplemented with NaCl powder to final concentration of 200mM, and with CHAPS powder to final concentration of 0.5% (w/v)). The cell suspension was put on ice for 10min with intermittent vortexing, and then centrifuged at 20,000g for 10min at 4C. The supernatant containing the AAV was collected. To set up the gravity column for AAV purification, 0.5ml of AAV8-binding slurry beads (ThermoFisher #A30789), or enough to purify AAV from up to eight 15cm dishes, was loaded into an empty column (Bio-Rad#731-1550). After the beads were tightly packed at the bottom, they were washed with 5ml of wash buffer (PBS supplemented with NaCl powder to a final concentration of 500mM). The supernatant containing AAV was then loaded onto the column. After all of the supernatant flowed through, the beads were washed with 10ml wash buffer twice. The AAV was then eluted with 3ml elution buffer (100mM glycine, 500mM NaCl in water, pH 2.5) and the eluate was immediately neutralized with 0.12ml 1M Tris-HCl (pH 7.5-8.0). The AAV was concentrated by centrifugation at 2000g for 3-5min at 4C using an 100k Amicon Ultra Centrifugal Filter Unit (Millipore #UFC810024). After centrifugation, the volume of AAV should be equal to or less than 0.5ml. The concentrated AAV was diluted with 4-5ml AAV dialysis buffer (PBS supplemented with powders to final concentrations of 212mM NaCl and 5% sorbitol (w/v)) and centrifuged at 2000g for 3-5min at 4C. The dilution and centrifugation processes were repeated 3 times. The final concentrated AAV was transferred into a 1.5ml tube and centrifuged at 20,000g for 5min to remove debris. The supernatant was aliquoted, flash frozen using liquid nitrogen, and stored at -80C.

### AAV injection

Six-week-old male mice were anaesthetized with isoflurane. About 0.5 × 10^11^ AAV particles were delivered into mice liver via orbital injection. Mice were allowed to recover for 2-3 weeks and then treated with HFD/WD for appropriate duration. AAV2/8 vectors were described as in Wang et al, PMID: 37040760.

### AAV induced liver specific *Pparg* KO mice

Liver-specific *Pparg* knockout mice were generated using an adeno-associated virus-based CRISPR system. AAV2/8 particles encoding by MOSAICS system, U6-driven single-guide RNA (sgRNA) targeting the mouse *Pparg* gene were constructed and packaged following above instructions. The sgRNA sequence (ATAAATAAGCTTCAATCGGA) was selected from a validated mouse gRNA sequences database (dbGuide database. https://frederick.cancer.gov/downloads/mm_guide_info.csv.gz). Eight-week-old Cas9 knock-in mice (Jackson#029415) were injected via orbital vein with 5 × 10^10^ viral genomes (vg) of AAV2/8-sg*Pparg* in 100 µL sterile PBS. Control mice received AAV2/8 expressing a non-targeting sgRNA. Mice were maintained for 2-3 weeks post-injection to allow for gene editing.

### H&E, immunofluorescence (IF), and Sirius Red staining

Liver pieces were fixed in buffered formalin (Fisherbrand #245-685) for 24h with gentle shaking at 4C and then transferred to 70%EtOH for another 24h with shaking at 4C. Paraffin embedding, liver sectioning (4mm thickness), and H&E staining were performed at the UT Southwestern Tissue Management Shared Resource Core. For Cryo-section IF staining, Tissue was collected immediately after euthanizing the mice and fixed with 4% paraformaldehyde overnight. The tissue was then washed with PBS 5 times, for 10 min each time and incubated in PBS with 30% sucrose for 8 h and then frozen in Tissue-Tek O.C.T. Compound (Sakura Finetek,4583). Frozen tissue was cut into 30-μm sections on a Leica CM3050 S cryostat. Sections were briefly rinsed with PBS and blocked with PBS/0.3% Triton X-100/5% FBS overnight. Sections were then stained with αSMA antibody (14395-1-AP, Proteintech, 1:100) or F4/80 antibody (28463-1-AP, Proteintech, 1:100) for 12 hours/overnight. Sections were washed with PBS/0.03% Triton X-100/5% FBS 5 times, for 1 h each time and then stained with anti-rabbit Alexa Fluor 488 (Thermo Fisher, A-11011, 1:500) for 2 hours. Sections were washed with PBS/0.3% Triton X-100/5% FBS 3 times, for 30 min each time and stained with BODYPI and DAPI for 30 min, wash with PBS/0.3% Triton X-100/5% FBS 3 times for 10 min each time and then mounted in ProLong Diamond Antifade Mountant (Thermo Fisher, P36965).

Sirius Red staining was performed on paraffin embedded liver sections using the Picro Sirius Red Staining Kit (Abcam #ab150681) according to the manufacturer’s protocol. Images were then taken on a ZEISS LSM 900 confocal microscope. ImageJ was used to quantify IF staining.

### Serum and liver metabolic assays

At the study endpoints, blood was collected immediately after sacrificing the mouse. Blood was let standing for 30 min and then centrifuged at 3000 g for 15 minutes at 4 °C. The supernatant (serum) was analyzed for AST, ALT, cholesterol, and triglyceride using specific reagent kits (VITROS 8433815, 1655281, 1669829, 1336544) on a fully automated OCD Vitros 350 dry chemistry analyzer. All analyses followed the protocols provided by the reagent kit manufacturer (Ortho Clinical Diagnostics, Raritan, NJ) at the UT Southwestern Metabolic Phenotyping Core. Or samples analyzed for AST, ALT, cholesterol, FFA and triglyceride using specific reagent kits (Cayman 701640, 700260, 10007640, 700310, 10010303) following the protocols provided by Cayman.

### Electron microscopy

For transmission electron microscopy (TEM), mice were first perfused transcardially with 15 mL of ice-cold fixative buffer (4% paraformaldehyde, 2.5% glutaraldehyde plus 0.2% picricc acid in 0.01M phosphate buffer) under deep anesthesia. Immediately after perfusion, small liver tissue blocks (∼1 mm³) were dissected and immersed in the same fixative buffer. The tissue was then post-fixed in 1% osmium tetroxide with 1.5 % K_3_[Fe(CN)_6_] in 0.1 M sodium cacodylate buffer for 1 h at room temperature, rinsed with water and *en bloc* stained with 0.5% aqueous uranyl acetate in 25% methanol overnight at 4 degrees C. After five rinses with water, specimens were stained with 0.02M lead nitrate in 0.03M L-aspartate for 30 minutes at 60 degrees C. Samples were dehydrated with increasing concentration of ethanol and infiltrated with Embed-812 resin and polymerized in a 60°C oven overnight. Images were acquired on a JOEL 1400 Plus transmission electron microscope equipped with a LaB6 source using a voltage of 120 kV, and the images were captured by an AMT BIOSPRINT 16M-ActiveVu mid mound CCD camera.

### Liver TG measurement

Liver and serum samples were collected from WT and *Clstn3b-*LKO cohorts or WT and *Clstn3b* LTG cohorts maintained on HFD/WD. About 60 mg of liver was removed from each mouse for measuring TG. TG levels were measured with the Triglyceride Colorimetric Assay Kit (Cayman, 10010303) or Glycerol Assay Kit (Sigma, MAK117) following the manufacturer’s instructions.

### Proteomics

LD was isolated from mouse liver without protease digestion. The suspension was mixed with 10x volume of acetone and incubated at -20℃ overnight for delipidation and protein precipitation. The mixture was centrifuged at 12,000 g for 5 min. The pellets were washed with acetone and dried by heating at 60℃ for at least 15 min. Pellets were then washed with 20% TCA to remove excess sucrose, washed with acetone, and dried. The pellets were dissolved in 50 mM triethylammonium bicarbonate (TEAB, pH=8), 5% SDS at 60℃ with shaking for 30 min. Protein concentration was determined with a BCA method.

Samples were then reduced by adding dithiothreitol (DTT) to a final concentration of 10 mM and samples were incubated at 56°C for 30 min. After cooling, iodoacetamide was added to a final concentration of 20 mM and samples were alkylated for 30 min at room temperature in the dark. Following centrifugation for 2 min at 13.2 krpm, the supernatants were removed and digested overnight with trypsin at 37°C using an S-Trap (Protifi). Following digestion, the peptide eluate was dried and reconstituted in 100 mM TEAB buffer.

Samples were injected onto an Orbitrap Fusion Lumos mass spectrometer coupled to an Ultimate 3000 RSLC-Nano liquid chromatography system (Thermo). Samples were injected onto a 75 um i.d., 75-cm long EasySpray column (Thermo) and eluted with a gradient from 0-28% buffer B over 180 min. Buffer A contained 2% (v/v) ACN and 0.1% formic acid in water, and buffer B contained 80% (v/v) ACN, 10% (v/v) trifluoroethanol, and 0.1% formic acid in water. The mass spectrometer operated in positive ion mode with a source voltage of 1.8-2.0 kV and an ion transfer tube temperature of 300°C. MS scans were acquired at 120,000 resolution in the Orbitrap and top speed mode was used for SPS-MS3 analysis with a cycle time of 2.5 s. MS2 was performed with CID with a collision energy of 35%. The top 10 fragments were selected for MS3 fragmentation using HCD, with a collision energy of 55%. Dynamic exclusion was set for 25 s after an ion was selected for fragmentation.

Raw MS data files were analyzed using Proteome Discoverer v2.4 SP1 (Thermo), with peptide identification performed using a trypsin digest search with Sequest HT (cleavage after Lys and Arg except when followed by Pro). The mouse reviewed protein database from UniProt (downloaded Jan. 28, 2022, 17,062 entries) was used. Fragment and precursor tolerances of 10 ppm and 0.6 Da were specified, and three missed cleavages were allowed. A minimum peptide length of 6 residues was required. Carbamidomethylation of Cys and TMT10plex labelling of N-termini and Lys sidechains were set as a fixed modification, with oxidation of Met set as a variable modification. The false-discovery rate (FDR) cutoff was 1% for all peptides. At least two unique peptides were required for protein identification.

### Liver LD isolation and digestion

The liver tissue was dissected, minced into tiny pieces with a spring scissor and transferred to a motorized homogenizer in HES buffer (20mM HEPES + 1mM EDTA + 250mM Sucrose). The homogenate was filtered with double-layer gauze and centrifuged at 2000 g for 5 min. The infranatant was removed with a syringe and the buoyant LD fraction was transferred with a wide-opening tip into 5 ml tubes and washed with HES buffer 2 times. The LD was then transferred to Ultra-Clear ultracentrifuge tubes (Beckman-Coulter), adjusted to a final concentration of 20% sucrose, and overlaid by 5% sucrose/HE and HE (20mM HEPES + 1mM EDTA). The gradient was centrifuged at 16,000 g for 10 min at 4℃. The buoyant LD was then collected.

To remove the LD-bound organelles, the LD fraction was digested with 1 mg/mL proteinase K for 5-15 min at 37℃ and 1mM PMSF was added to inactivate proteinase K. The mixture was centrifuged through the a before mentioned sucrose gradient at 210,000g for 1 hr. The buoyant LD was then collected.

### Phospholipids quantitation

LD was isolated from mouse liver and digested with proteinase K following the procedure described above. The collected LD fractions (100-200 μL suspension in HES buffer) were transferred into round bottom glass tubes and extracted with a mixture of 1 mL hexane, 1 mL methyl acetate, 0.75 mL acetonitrile and 1 mL water as previously described. The extracts were vortexed for 5 s and centrifuge at 2,671 g for 5 min to partition into 3 phases. The upper and middle phases were collected into separate glass tubes and dried under N_2_. Phospholipids in the middle phase were measured with the Phosphatidylcholine Assay Kit (Colorimetric/Fluorometric) (Abcam, ab83377) and Phosphatidylethanolamine Assay Kit (Fluorometric) (Sigma, MAK361) following the manufacturer’s instructions. TG in the upper phase was dissolved in 350 μL ethanolic KOH (2 part EtOH and 1 part 30% KOH) and incubated overnight at 55°C for complete hydrolysis. The volume was then brought to 1200 μL with H2O: EtOH (1:1) and vortexed to mix. Two hundred μL was transferred to a new tube, mixed with 215 μL 1M MgCl_2_ and vortexed. The mixture was incubated on ice for 10 min and centrifuged at 13,000g for 5 min. Glycerol content in the supernatant was determined with the Free Glycerol Reagent (Sigma, F6428) following the manufacturer’s instructions.

To calculate LD surface phospholipids density, we normalized phospholipids abundance as measured by the fluorometric kit or phospholipidomics to LD surface area of each sample. The detailed normalization procedure is described in a previous publication (Zhang et al, PMID: 38293096). Briefly, we divided total TG content of each sample by the mean LD volume to derive total LD number, which was then multiplied by the mean LD surface area. Mean LD volume and surface area were determined by Imaris analysis of LD images harvested from the same batch of sample.

#### Primary hepatocyte isolation and culture

Primary hepatocytes were isolated from mouse livers using a two-step collagenase perfusion method. Briefly, mice were anesthetized with isoflurane, and the abdominal area was sterilized with 70% ethanol. A midline incision was made to expose the portal vein, and a catheter was inserted into the inferior vena cava (IVC) and secured by threading a needle through the catheter’s butterfly wing. Perfusion was initiated with Liver Perfusion Medium (Life Technologies, #17701038) at a flow rate of 3 mL/min, using approximately 30–50 mL of medium. The portal vein was immediately severed to allow for adequate outflow. Following perfusion with Liver Perfusion Medium, the liver was perfused with Liver Digestion Medium (Life Technologies, #17703034) at the same flow rate, using an additional 30–50 mL. After digestion, the liver was carefully removed and placed in 10 mL of Liver Digestion Medium in a 10 cm culture dish. The liver capsule was gently peeled away, and the tissue was gently agitated to release hepatocytes. The cell suspension was then filtered through a 70 μm nylon mesh into a 50 mL tube prefilled with 20 mL of Hepatocyte Washing Medium (Life Technologies #17704024) on ice to halt enzymatic activity. Cells were pelleted by centrifugation at 50 g for 5 minutes at 4°C, and the supernatant was discarded. The cell pellet was resuspended in 20 mL of Washing Medium, and the wash step was repeated twice to ensure cell purity. Cell viability and yield were assessed by Trypan blue exclusion. Cells were then centrifuged once more at 50 g for 5 minutes and resuspended in the desired volume of washing medium or buffer for further experiments.

### Primary hepatocyte Lipolysis assay

After isolation, primary hepatocytes were seeded onto collagen-coated 6-well plates. To prevent mitochondrial oxidation or re-esterification of free fatty acids (FFAs) released via lipolysis, an ACSL inhibitor (10uM Triacsin C, Cat#2472, Biotechne) and a CPT1 inhibitor (10uM Etomoxir, Cat#HY-50202, MCE) were added upon switching to serum-free medium. After 24 hours of incubation, the culture medium was collected, and the FFA concentration was subsequently measured with the Free fatty acid fluorometric assay kit (Cayman, 700310) following the manufacturer’s instructions. To evaluate ATGL and HSL functional relevance, hepatocytes were treated with inhibitors of ATGL (20uM Atglistatin, Cat# HY-15859, MCE) or HSL (2uM HSL-IN-1, Cat# SML3763, Sigma) for 12 hours in vitro and then took the fixed cell imaging.

### Fixed cell imaging

Hepatocytes were isolated as above described and attached to glass coverslips pre-coated with laminin and poly-D-lysine. Images were taken on a ZEISS 900 confocal microscope with Airyscan.

### Image analysis

To analyze isolated lipid droplets, images were imported to Imaris 10.2.0 (Bitplane). Individual LDs were rendered by the “Surface” function and surface area and volume were measured. The rendering parameters were kept the same between treatments in each experiment. To analyze the extent of CLSTN3B wrapping of LDs, individual LDs and the surrounding CLSTN3B signals were rendered by the “Surface” function. The fraction of the volume bound by the LD surface overlapping with the volume bound by the CLSTN3B signal surface was calculated with the “Object-Object statistics” function. All plots were generated in R.

Immunofluorescence staining for aSMA and F4/80 was performed on frozen liver sections, and fluorescence signal was analyzed using ImageJ (NIH). Images were captured under identical exposure settings using ZEISS 900 confocal microscope. Red fluorescence signals (representing aSMA or F4/80) were extracted by splitting color channels, and a threshold was applied to define positive signal areas. The percentage of aSMAL or F4/80L area was calculated as the ratio of red-positive area to total tissue area. All signal analyses were performed in ImageJ.

### Hepatocyte respiration

Hepatocyte oxygen consumption data were collected by high-resolution respirometry with an Oroboros Oxygraph-2k (Oroboros Instruments, Innsbruck, Austria) in a standard configuration, with 2 ml volume of the two chambers, at 37oC, and 350 rpm stirrer speed. Specifically, freshly Isolated primary hepatocytes were suspended at a density of 1–2 ×10^6^ cells/mL in a respiration buffer consisting of 100mM potassium chloride, 2mM magnesium chloride, 1mM EDTA, 4mM dibasic potassium phosphate, 20mM HEPES, and 0.1% (w/v) BSA at pH 7.2, and same number of cells were added to the respirometry chamber for measurement. The software DatLab7 was used for data acquisition (2s time intervals) and analysis. All measurements were performed within 4 hours of cell isolation.

### Anion-Exchange Chromatography for ATP and ADP measurement in hepatocytes

Primary hepatocytes cultured in 12 well plates were extracted with 500ul solvent containing Methanol:Acetonitrile:Water 2:2:1 (v/v/v). The extract was transferred to a 1.5ml Eppendorf tube and subjected to three freeze-thaw cycles between liquid nitrogen and 37°C water bath. After the third thaw, the samples were vortexed for 1 min and then centrifuged at a maximum speed for 15 minutes in a refrigerated centrifuge. The supernatant was transferred to a new tube and then dried in a SpeedVac to a volume just below 100ul. The sample was diluted to 500ul by adding 5 mM Tris-HCL 8.0, and resolved on Capto HiRes Q 5/50 0-40%B in 15ml (Buffer A: 5 mM Tris-HCl 8.0; Buffer B: 5mM Tris-HCl 8.0 and 1M NaCl). The ATP and ADP peaks were monitored and quantified based on UV absorption at 254 nm.

### Real-time qPCR analysis

The following primers were used for qPCR analysis of gene expression.

*Clstn3b*-fwd, CTCCGCAGGAACAGCAGCCC, rev, AGGATAACCATAAGCACCAG; *tnf*-fwd, GGTGCCTATGTCTCAGCCTCTT, rev, GCCATAGAACTGATGAGAGGGAG; *ccl2*-fwd, GCTACAAGAGGATCACCAGCAG, rev, GTCTGGACCCATTCCTTCTTGG; *adgre1*-fwd, CGTGTTGTTGGTGGCACTGTGA, rev, CCACATCAGTGTTCCAGGAGAC; *lipe*-fwd, TTGGGGAGCTCCAGTCGGA, rev, TCGTGCGTAAATCCATGCTGT; *fabp4*-fwd, ACACCGAGATTTCCTTCAAACTG, rev, CCATCTAGGGTTATGATGCTCTTCA; *L19*-fwd, GGTCTGGTTGGATCCCAATG, rev, CCCATCCTTGATCAGCTTCCT; *lpl*-fwd, GGGAGTTTGGCTCCAGAGTTT, rev, TGTGTCTTCAGGGGTCCTTAG; *cfd*-fwd, CATGCTCGGCCCTACATGG, rev, CACAGAGTCGTCATCCGTCAC; pparg-fwd, GTACTGTCGGTTTCAGAAGTGCC, rev, ATCTCCGCCAACAGCTTCTCCT; ppara-fwd, AGAGCCCCATCTGTCCTCTC, rev ACTGGTAGTCTGCAAAACCAAA; ppargc1a-fwd, GAATCAAGCCACTACAGACACCG, rev, CATCCCTCTTGAGCCTTTCGTG; cpt2-fwd, CAGCACAGCATCGTACCCA, rev,TCCCAATGCCGTTCTCAAAAT cpt1a-fwd, CTCCGCCTGAGCCATGAAG, rev,CACCAGTGATGATGCCATTCT; acads-fwd, TGGCGACGGTTACACACTG, rev,GTAGGCCAGGTAATCCAAGCC; acadl-fwd, TCTTTTCCTCGGAGCATGACA, rev, GACCTCTCTACTCACTTCTCCAG; acadm-fwd, AGGGTTTAGTTTTGAGTTGACGG, rev, CCCCGCTTTTGTCATATTCCG; acaa2-fwd, GAATCAAGCCACTACAGACACCG,rev, CATCCCTCTTGAGCCTTTCGTG; etfa-fwd, GTCTTGGAGGTGAAGTGTCCTG, rev, GCATCATGCTGAGCCACCAGAA ; etfdh-fwd, ATGGAGGCTCTTTCCTTTACCAC rev, GCTTACACCTCTGGAACTCTCG ; echs1-fwd, CTCAACCAAGCACTGGAGACCT rev, GCTGGAGTAACAGTCCTGAAATG ; hadhb-fwd, CACTGCGTTCTCATAGTCTGGC rev, GCCATTTGCTCCAGTGAGGAAG; hadha-fwd, GTTTGAGGACCTCGGTGTAAAGC rev, GAGAGCAGATGTGTTGCTGGCA; tgfb1-rwd, TGATACGCCTGAGTGGCTGTCT, rev, CACAAGAGCAGTGAGCGCTGAA; timp1-fwd, TCTTGGTTCCCTGGCGTACTCT rev, GTGAGTGTCACTCTCCAGTTTGC; acat2-fwd, GAGATTGTGCCAGTGCTGGTGT, rev, GTGACAGTTCCTGTCCCATCAG ; cola1-fwd, CCTCAGGGTATTGCTGGACAAC, rev, CAGAAGGACCTTGTTTGCCAGG ; mmp2-fwd, CAAGGATGGACTCCTGGCACAT, rev, TACTCGCCATCAGCGTTCCCAT; mmp9-fwd, GCTGACTACGATAAGGACGGCA, rev, TAGTGGTGCAGGCAGAGTAGGA; desmin-fwd, GCGGCTAAGAACATCTCTGAGG, rev, ATCTCGCAGGTGTAGGACTGGA; pdgfrb-fwd, GTGGTCCTTACCGTCATCTCTC, rev, GTGGAGTCGTAAGGCAACTGCA; tnfa-fwd, GGTGCCTATGTCTCAGCCTCTT, rev, GCCATAGAACTGATGAGAGGGAG; bax-fwd, AGGATGCGTCCACCAAGAAGCT, rev, TCCGTGTCCACGTCAGCAATCA. actin-fwd, CATTGCTGACAGGATGCAGAAGG, rev, TGCTGGAAGGTGGACAGTGAGG.

### ChIP-qPCR

The liver tissue was dissected, minced into tiny pieces with a spring scissor and transferred to a motorized homogenizer in HES buffer (20mM HEPES + 1mM EDTA + 250mM Sucrose). Add 32% PFA into homogenizing mixture to final 1% concentration and crosslink by gently rocking at room temperature for 10 minutes, followed by quenching with 0.125 M glycine. Nuclei were collected by centrifugation and lysed with nuclei lysis buffer (1% SDS, 50 mM Tris-HCl, 10 mM EDTA at pH 8.0) and sonicated to shear DNA to ∼100–500 bp fragments. The lysate was diluted with a dilution buffer (0.11% sodium deoxycholate, 1.1% Triton X-100, 50 mM Tris-HCl, 167 mM NaCl at pH 8.0) and kept some lysate to yield input DNA. Incubate the diluted lysate with anti-PPARγ antibody (Proteintech, Cat#6643-1-AP) overnight at 4°C. The protein–DNA complex was isolated with protein G Dynabeads, washed, and eluted off the beads by elution buffer (0.5% SDS, 10 mM Tris-HCl pH8, 5 mM EDTA at pH 8.0). The eluent was treated with RNase A and proteinase K before reversing crosslinking. DNA was then harvested by ethanol precipitation and analyzed by qPCR using primers targeting PPARγ-binding regions and non-binding regions in selected gene promoters. Input was used as control.

### Metabolic cage

Energy expenditure and metabolic parameters were assessed using a comprehensive lab animal monitoring system (CLAMS; Columbus Instruments) and operated by the UT Southwestern Metabolic Phenotyping Core. Mice were singly housed in metabolic cages for continuous monitoring of oxygen consumption (VO₂), carbon dioxide production (VCO₂), respiratory exchange ratio (RER), locomotor activity, food intake, and water consumption. Animals were acclimated in the chambers for 24 hours prior to data collection. Measurements were recorded over a 45-hour period under a 12-hour light/dark cycle at 30°C with ad libitum access to food and water. Data were analyzed using the manufacturer’s software and normalized to body weight.

### Temperature measurement

Core body temperature of mice was measured using a rectal thermometer (BAT-12, PHYSITEMP) with a lubricated probe inserted ∼2 cm into the rectum. Mice were housed in climate Chamber at 30L. Mice were gently restrained by hand, and temperature was recorded once a stable reading was obtained, typically within 10–15 seconds. All measurements were performed at the same time of day at midnight. To assess surface temperature over specific anatomical regions by infrared thermograph, mice were anesthetized briefly, and hair was removed from the skin overlying the liver (right upper abdominal area) and interscapular brown adipose tissue (BAT) using hair Clipper and the treat with commercial depilatory cream. After full recovery from anesthesia, mice were placed individually in a temperature-controlled environment, and thermal images were captured using an infrared camera (TC004, TOPDON TECH). Surface temperatures over the liver and BAT regions were quantified using ImageJ to measure the signal intensity and compared with the temperature bar to calculate the corresponding temperature.

### FAO blue staining

Fatty acid oxidation (FAO) activity was assessed using the FAO Blue™ kit (FNK-FDV-0033, Diagnocine) according to the manufacturer’s instructions. Briefly, cells were incubated with the 20uM FAO Blue working solution at 37°C for 30 minutes in the dark. After staining, cells were washed with the PBS and changed to fresh medium without serum and subjected to live-cell imaging using a fluorescence microscope (excitation/emission: 405/450 nm).

### TMRE staining

Mitochondrial membrane potential staining was performed using TMRE (I34361, Thermo Fisher) following the product instruction. Briefly, cells were incubated with 100 nM TMRE diluted in culture medium at 37°C for 30 minutes. After incubation, cells were immediately subjected to live-cell imaging using a fluorescence microscope (excitation/emission: 548/574 nm) without washing.

### LD size distribution plotting

To visualize the relative volume distribution of LDs, a volume-weighted density graph was plotted in which the x value is LD volume (log transformed) and the y value is the density weighted by the relative percentage of total LD volume. To characterize the volume-weighted volume distribution, a probability function was first generated from the density distribution by the “approxfun” function in R. The median value of the distribution was then calculated by random simulation (“simulate” function, 1e6 times) of the probability function in R.

### MASLD patient cohorts

Association of hepatic CLSTN3B expression and clinical phenotypes/outcomes was assessed in two previously published cohorts of MASLD patients (Govaere et al, PMID: 33268509; Fujiwara et al, PMID: 35731891). The first cohort includes 206 European Caucasian patients with MASLD at various fibrosis stages and 10 healthy obese controls, for which hepatic transcriptome dataset is available (NCBI Gene Expression Omnibus, accession number GSE135251). The second cohort includes 106 non-cirrhotic Japanese patients with MASLD, among which 77 patients underwent paired liver biopsies with a median interval of 2.3 years and transcriptome profiling (GSE193084).

### Quantitation of *CLSTN3B* gene expression level

Raw sequencing reads in FASTQ format were preprocessed using Cutadapt (v2.5) (Martin, M. Cutadapt removes adapter sequences from high-throughput sequencing reads. EMBnet.journal 17, 10–12 (2011)) to trim low-quality bases and adapter sequences (parameters “-q 20 --max-n 3 -a Ill_Univ_Adapt”). The trimmed reads were then aligned to the human reference genome (GRCh38.p14) using STAR aligner (v2.7.3a) (Dobin et al, PMID: 23104886) with parameters “--chimSegmentMin 15 --chimJunctionOverhangMin 15”. Gene-level quantification was performed using featureCounts (v1.6.3) (Liao et al, PMID: 24227677), assigning aligned reads to annotated gene features. Expression levels of CLSTN3B and CLSTN3 genes were quantified using exons unique to each transcript based on ENST00000535313.2 from the GENCODE release 45 annotation. Raw read counts were normalized using the relative log expression (RLE) method implemented in the DESeq2 R package (v1.42.0) (Love et al, PMID: 25516281). Bioinformatic data analysis was performed using BioHPC supercomputing facility at University of Texas Southwestern Medical Center.

### Statistics and Reproducibility

All data shown are mean ± s.e.m. Statistical significance was calculated by unpaired Student’s two-sided t-test for comparisons between two groups and one-way ANOVA with Tukey’s post hoc test for comparisons between three groups. Two-way Repeated Measurement ANOVA was used to analyze time-series data. To assess statistical differences between pair-wise comparisons in LD size analysis, the sample size, mean and standard error of each comparison sample was estimated from Two-way ANOVA with Tukey’s correction. A second one-way ANOVA with Tukey’s correction was then applied to compare the difference between pair-wise comparison samples. When analyzing the impact of lipase inhibitors on LD size, an arbitrary LD volume cutoff of 1 μm^3^ was applied to mitigate the confounding effect of a large number of small LDs resulting from re-esterification on the enlargement of a relatively small number of large LDs induced by lipase inhibitors. All experiments have been successfully repeated with similar results at least three times.

## Acknowledgement

We thank Dr. Xun Wang at Children’s Research Institute at UTSW for the guidance on primary hepatocyte isolation from mice on HFD; Andrew Lemoff and the Proteomics core at UTSW for the proteomics analysis; the Molecular Pathology core at UTSW for the histology analysis; the EM core at UTSW (supported by NIH grant 1S10OD021685-01A1) for project discussion and EM sample preparation. The automated bioinformatics pipeline was developed by J.C. of the Data Science Shared Resource (DSSR) at UTSW Simmons Comprehensive Cancer Center (SCCC). Bioinformatics analyses were conducted on BioHPC, High-Performance Computing (HPC) resources embedded in Lyda Hill Department of Bioinformatics at UTSW. X.Z. is a Rita C. and William P. Clements, Jr. Scholar in Biomedical Research. This study was supported by the Endowed Scholars in Medical Science Program at UTSW, Cancer Prevention and Research Institute of Texas grant RR200084, NIH R01DK135556, and American Heart Association Award 23CDA1050474 to X.Z. Y.H. is supported by U.S. National Institutes of Health (CA233794, CA255621, CA282178, CA288375, CA283935); European Commission (ERC-AdG-2020-101021417); Cancer Prevention and Research Institute of Texas (RR180016, RP200554). H.S. is supported by Uehara Memorial Foundation. J.C. and JL are partly supported by the UTSW Simmons Comprehensive Cancer Center (SCCC) grant P30CA142543.

## Author contributions

C.Z and X.Z conceived the project. C.Z. and X.Z. designed the experiments. C.Z., D.Y., M.L., and X.Z. performed the bulk of the experiments. H.S., J.C., J.L., and Y.H. analyzed the association between human *CLSTN3B*/*CLSTN3* expression and MASLD parameters. J.W., Q.Z., and H.Z. provided the MOSAICS system for AAV-mediated gene deletion in hepatocytes, prepared AAV for exogenous gene expression in the liver, and provided guidance on establishing primary hepatocyte cultures. M.Y. performed image and statistical analyses. J.Z. performed liver cryosections and immunostaining for fibrosis analysis. X.Z and C.Z wrote the manuscript. All authors participated in reviewing and discussing the manuscript.

---

**Figure S1.**
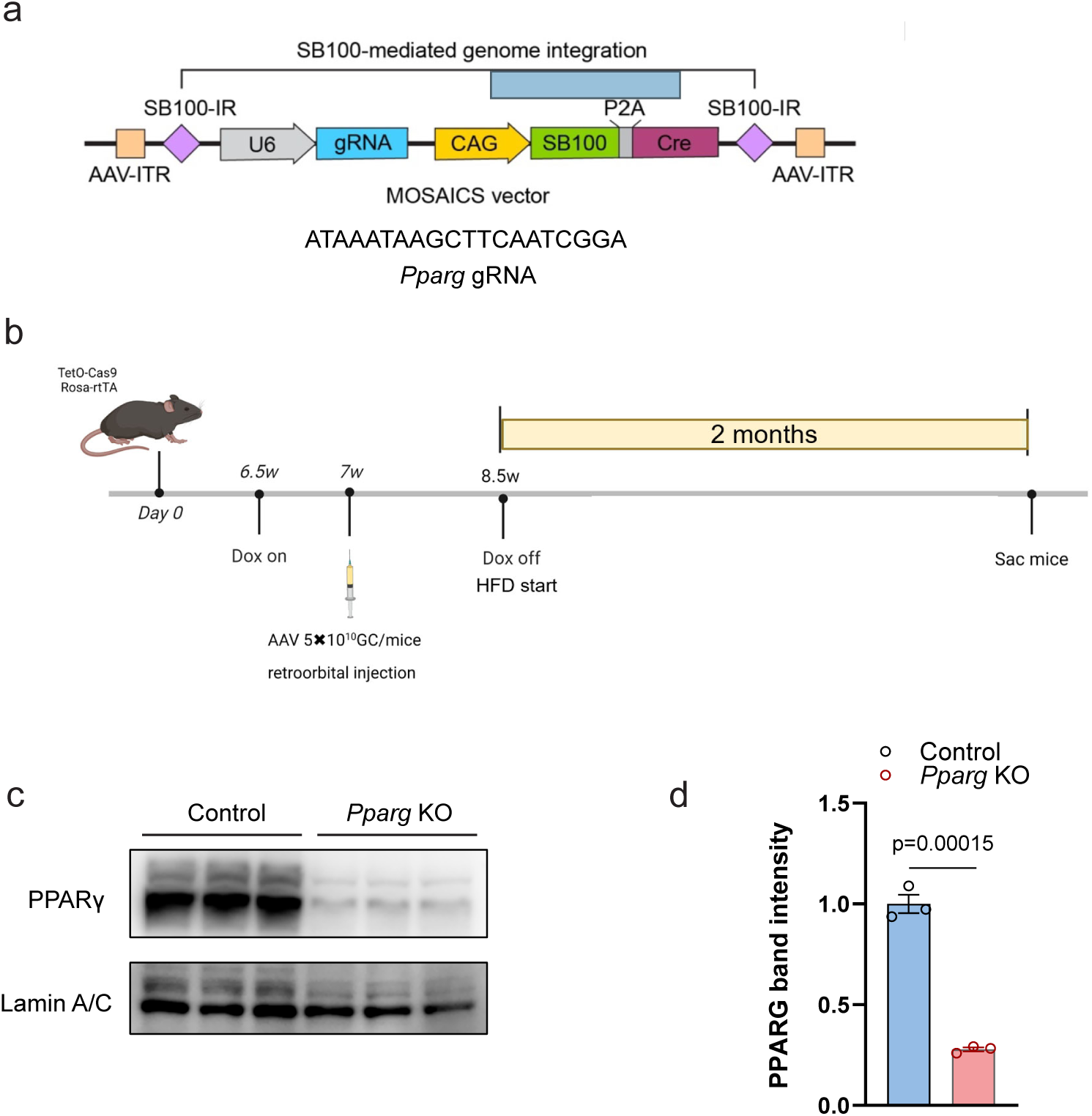
**a**, Schematic diagram of the MOSAICS vector containing *Pparg* gRNA for packaging adenovirus. **b**, *Pparg* KO mouse construction and HFD induction timeline. **c**, western blot analysis of PPARγ expression in nuclear extracts isolated from liver of control or *Pparg* KO mice on HFD (n=3). Data are mean ± s.e.m. Statistical significance was calculated by unpaired Student’s two-sided t-test in **d**.

**Figure S2.**
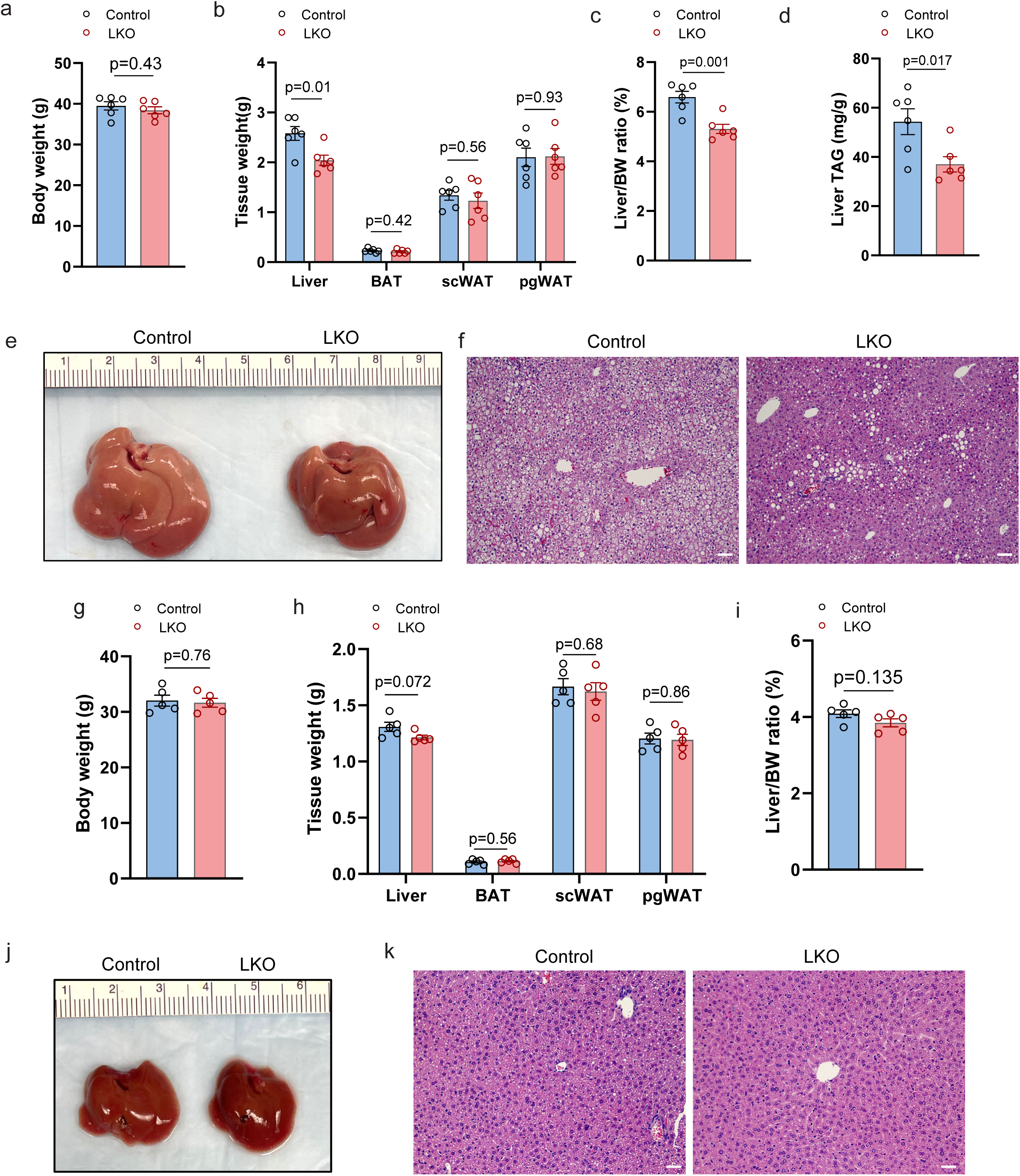
**a**-**f**, Body weight (**a**), tissue weights (**b**), liver/body weight ratio (**c**), liver TAG content (**d**), liver gross appearance (**e**) and histology (**f**) from WT control and LKO mice on WD for 10 weeks (n=5). **g**-**k**, Body weight (**g**), tissue weights (**h**), liver/body weight ratio (**i**), liver gross appearance (**j**) and histology (**k**) from WT control and LKO female mice on HFD for 10 weeks (n=5). Scale bar: 100 μm in **f** and **k**. Data are mean ± s.e.m. Statistical significance was calculated by unpaired Student’s two-sided t-test in all panels.

**Figure S3.**
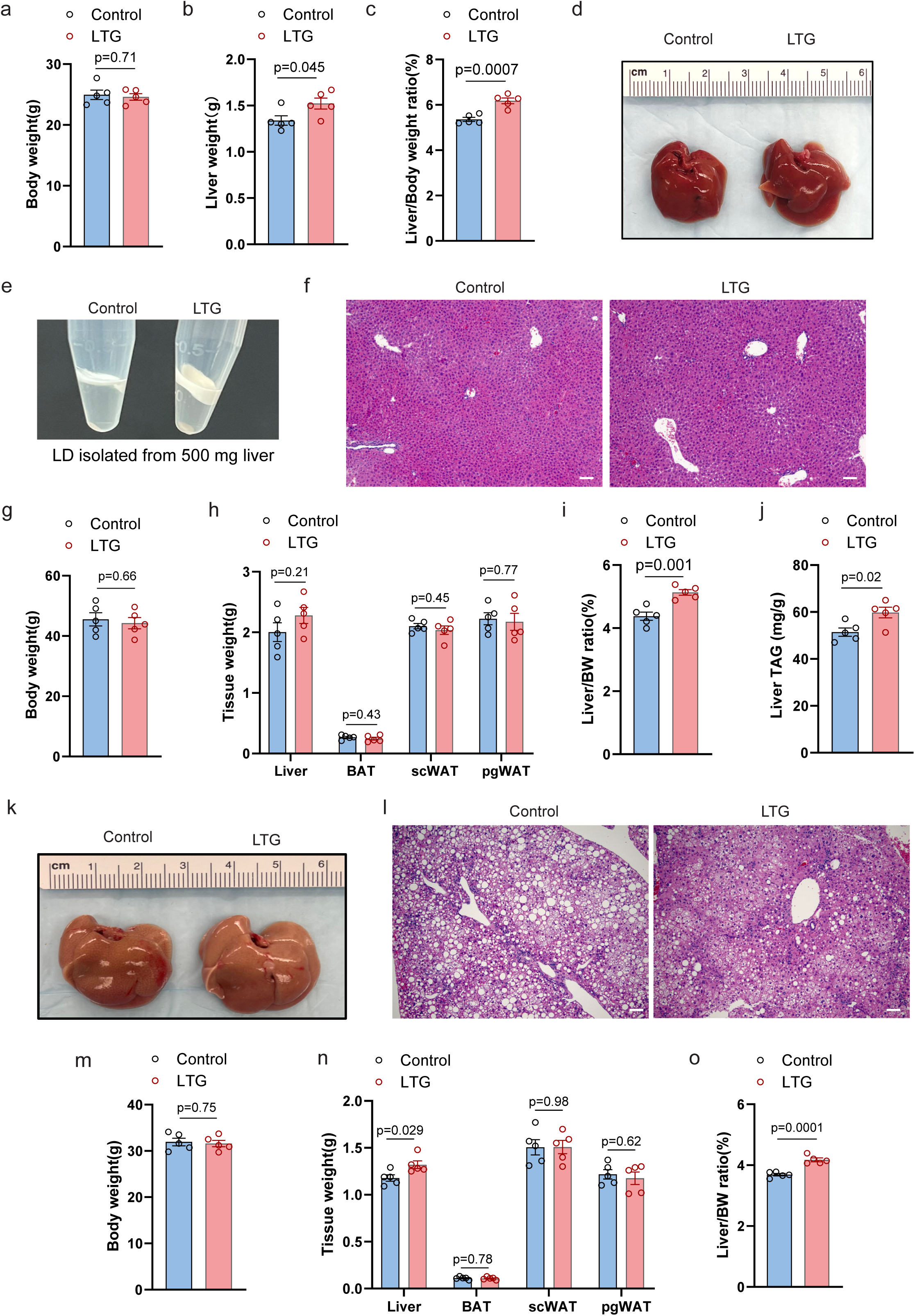
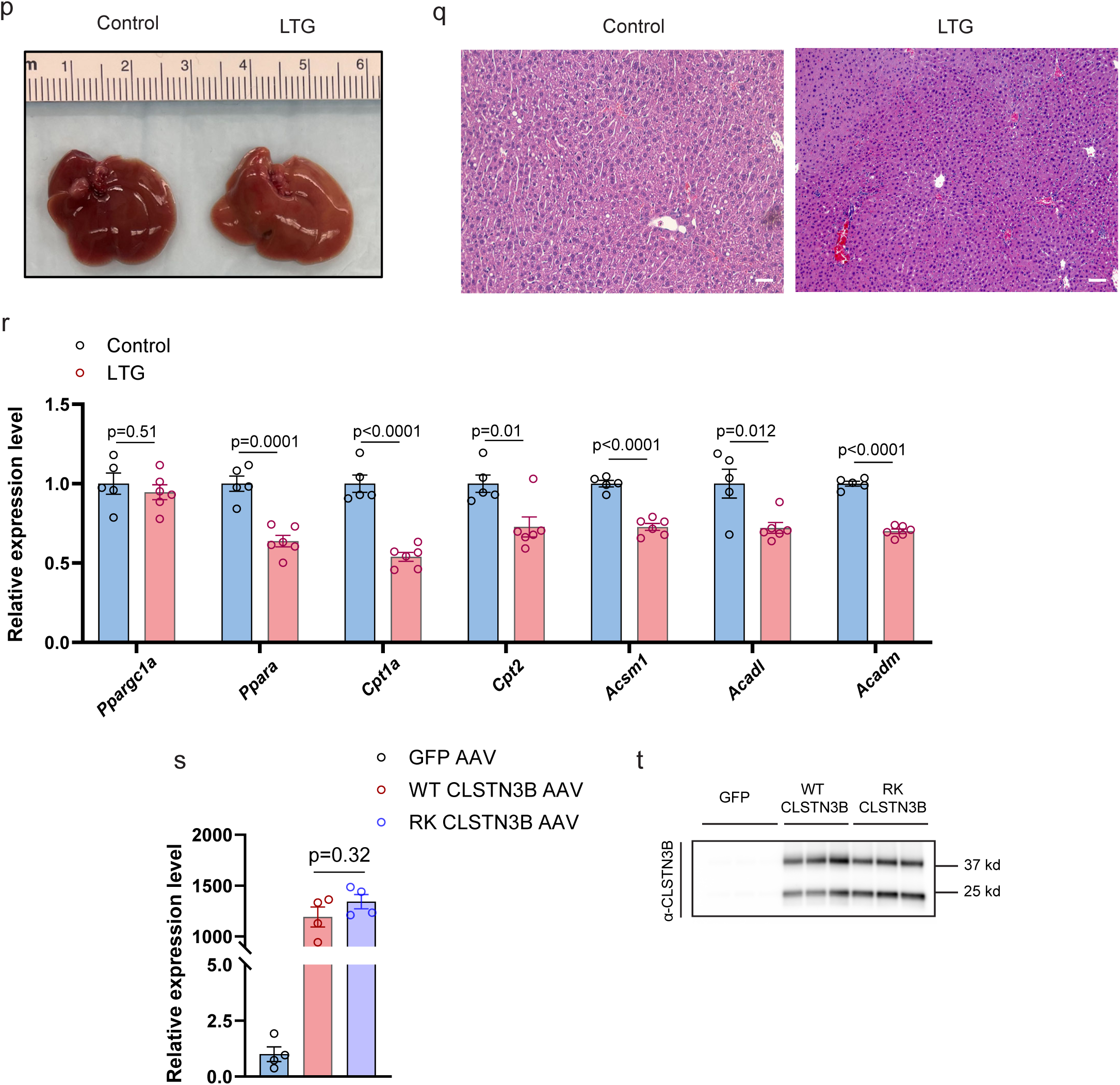
**a**-**f**, Body weight (**a**), liver weight (**b**), liver/body weight ratio (**c**), liver gross appearance (**d**), histology (**e**) and LD layer isolated from 500mg liver tissue (**f**) from male WT control and LTG mice on chow diet (n=5). **g**-**l**, Body weight (**g**), tissue weights (**h**), liver/body weight ratio (**i**), liver TAG content (**j**), liver gross appearance (**k**), and histology (**l**) from male WT control and LTG mice on WD for 10 weeks (n=5). **m**-**r**, Body weight (**m**), tissue weight (**n**), liver/body weight ratio (**o**), liver gross appearance (**p**) and histology (**q**), qPCR analysis of hepatic FAO gene expression (**r**) from WT control and LTG female mice on HFD for 10 weeks (n=5). (**s**), *Clstn3b* mRNA expression level in liver from mice receiving AAV-GFP, AAV-WT-CLSTN3B or AAV-RK-CLSTN3B injection. (**t**), CLSTN3B protein level on liver LD from mice receiving AAV-GFP, AAV-WT-CLSTN3B or AAV-RK-CLSTN3B injection. Scale bar: 100 μm in **f**, **l**, and **q**; Data are mean ± s.e.m. Statistical significance was calculated by unpaired Student’s two-sided t-test in **a**, **b**, **c**, **g**, **h**, **i**, **j**, **m**, **n**, **o**, and **r**, and One-way ANOVA with Tukey’s post hoc test in **s**.

**Figure S4.**
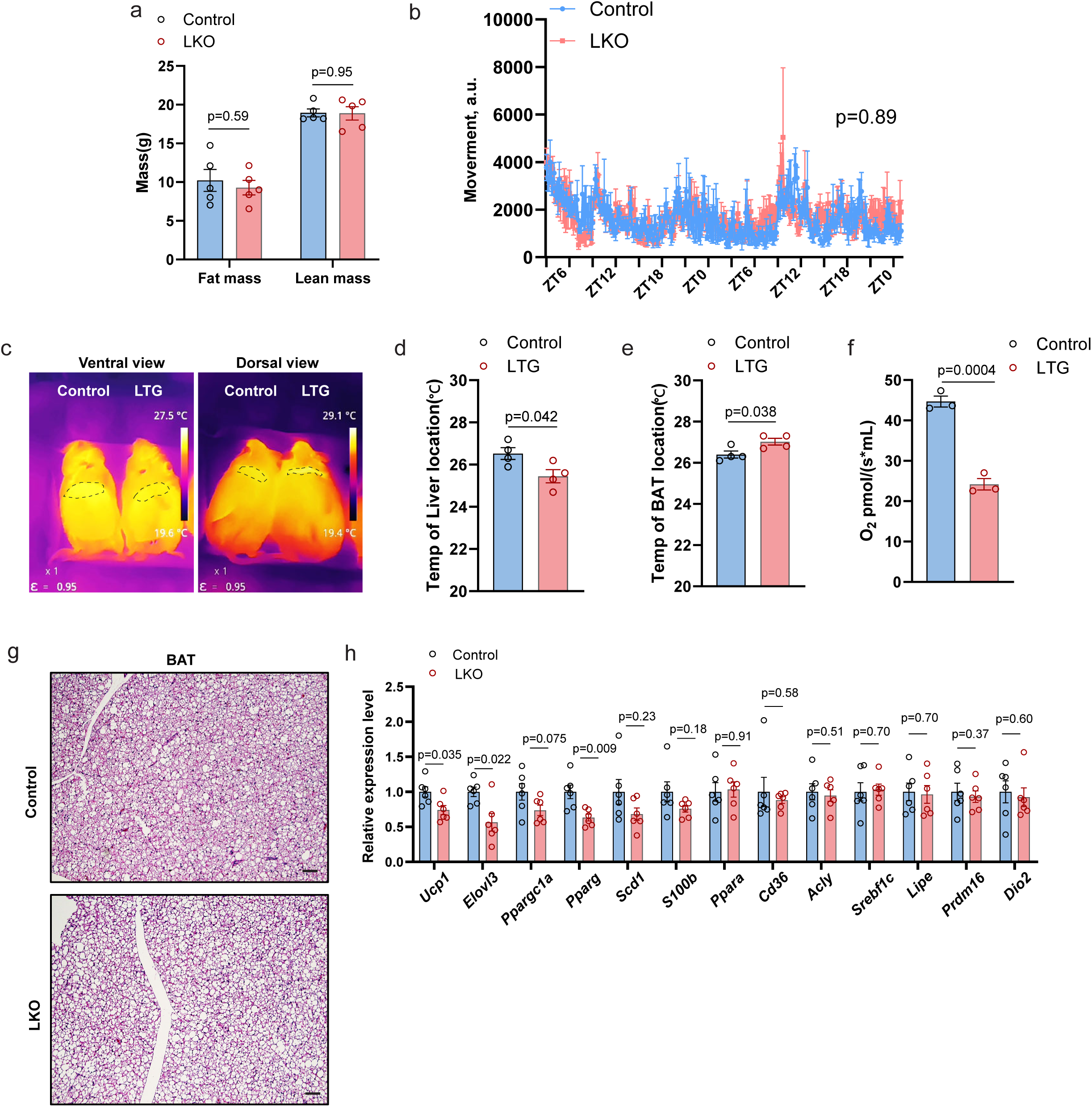
**a**, Fat mass and lean mass of WT control and LKO mice (n=5). **b**, Movement count of a WT control and LKO cohort over a 45-hour period (n=5). **c**-**e**, Infrared thermography images (**c**) and quantitation of surface temperature in the liver (**d**) and BAT region (**e**) of WT and LTG mice (n=4). **f**, Quantitation of O2 consumption rates from respiration assay of freshly isolated primary hepatocytes from WT and LTG mice (n=3 biological replicates). **g**-**h**, histology (**g**) and qPCR analysis of gene expression (**h**) of BAT from WT control and LKO mice maintained on HFD (n=5). Scale bar: 100 μm in **g**. Data are mean ± s.e.m. Statistical significance was calculated by unpaired Student’s two-sided t-test in **a**, **d**, **e**, **f**, and **h**, and repeated measurement two-way ANOVA in **b**.

**Figure S5.**
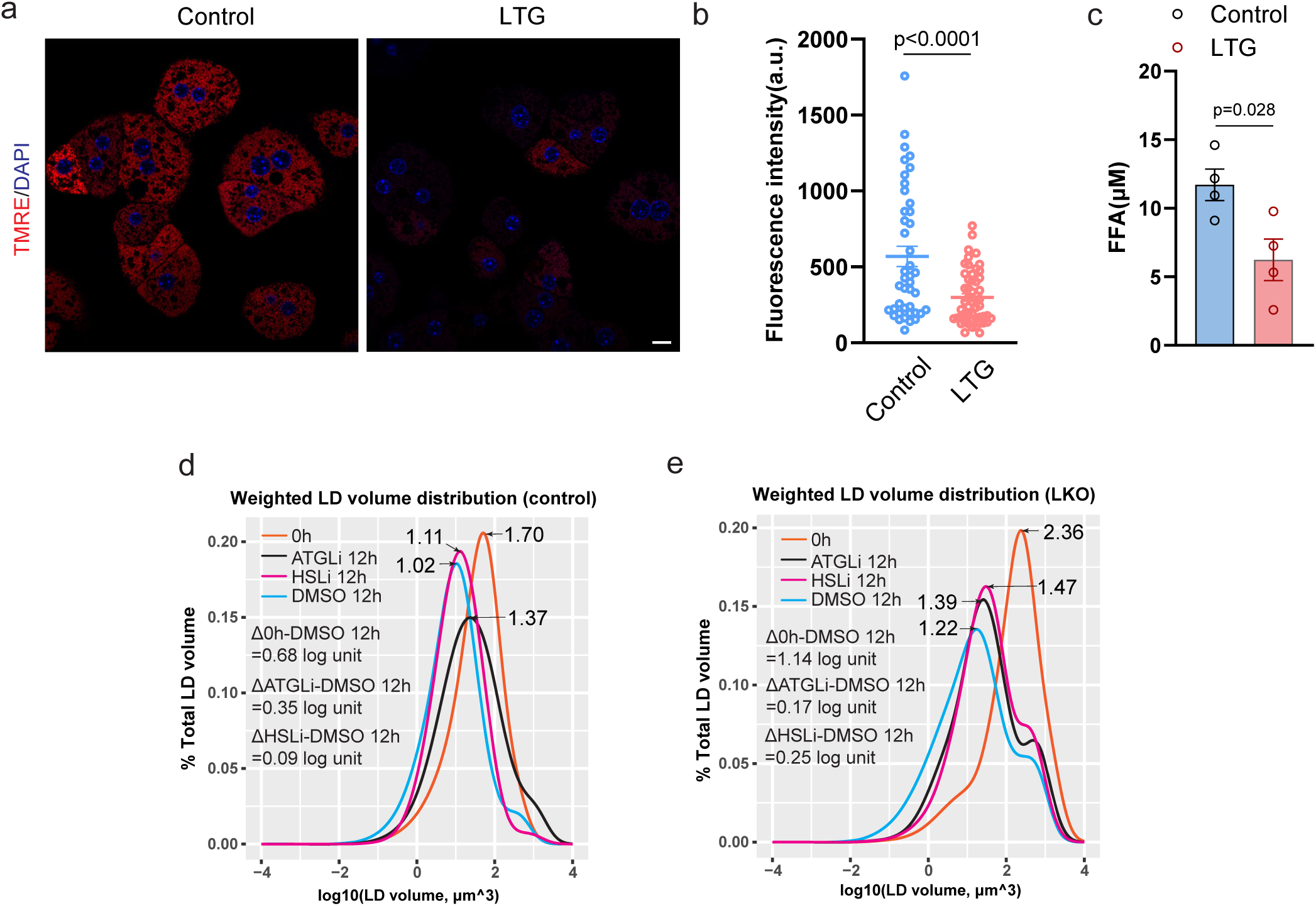
**a**, Lipolysis assay of WT and LTG primary hepatocytes (n=4 biological replicates). **b**-**c**, Fluorescence image (**b**) and quantitation (**c**) of mitochondrial membrane potential measurement by TMRE staining in WT and LTG primary hepatocytes (n=31-38). **d**-**e**, weighted LD volume distribution curve of WT (**d**) and LKO (**e**) primary hepatocytes treated with DMSO or lipase inhibitors. The shift in the peak value between conditions is indicated in the graph. Scale bar=5 μm. Statistical significance was calculated by unpaired Student’s two-sided t-test.

**Figure S6.**
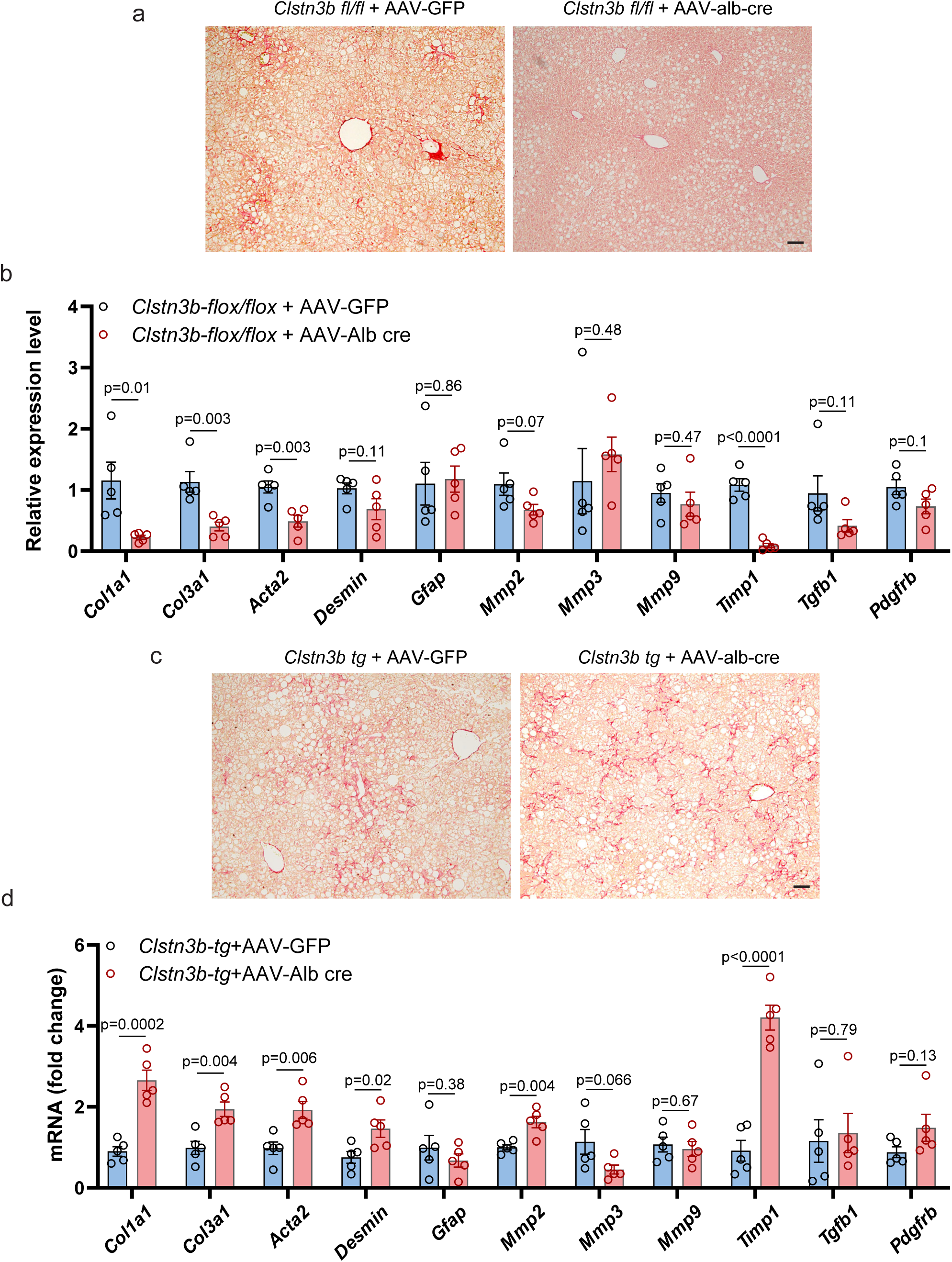
**a**, Sirium Red staining of liver sections of control and AAV-induced liver-specific CLSTN3B KO cohort (n=5) maintained on a WD. **b**, qPCR analysis of genes associated with hepatic fibrosis in the liver of mice (n=5) from **a**. **c**, Sirium Red staining of liver sections of control and AAV-induced liver-specific CLSTN3B transgenic cohort (n=5) maintained on a WD. **d**, qPCR analysis of genes associated with hepatic fibrosis in the liver of mice (n=5) from **c**. Scale bar: 100 μm in **a** and **c**. Data are mean ± s.e.m. Statistical significance was calculated by unpaired Student’s two-sided t-test in all panels.

**Figure S7.**
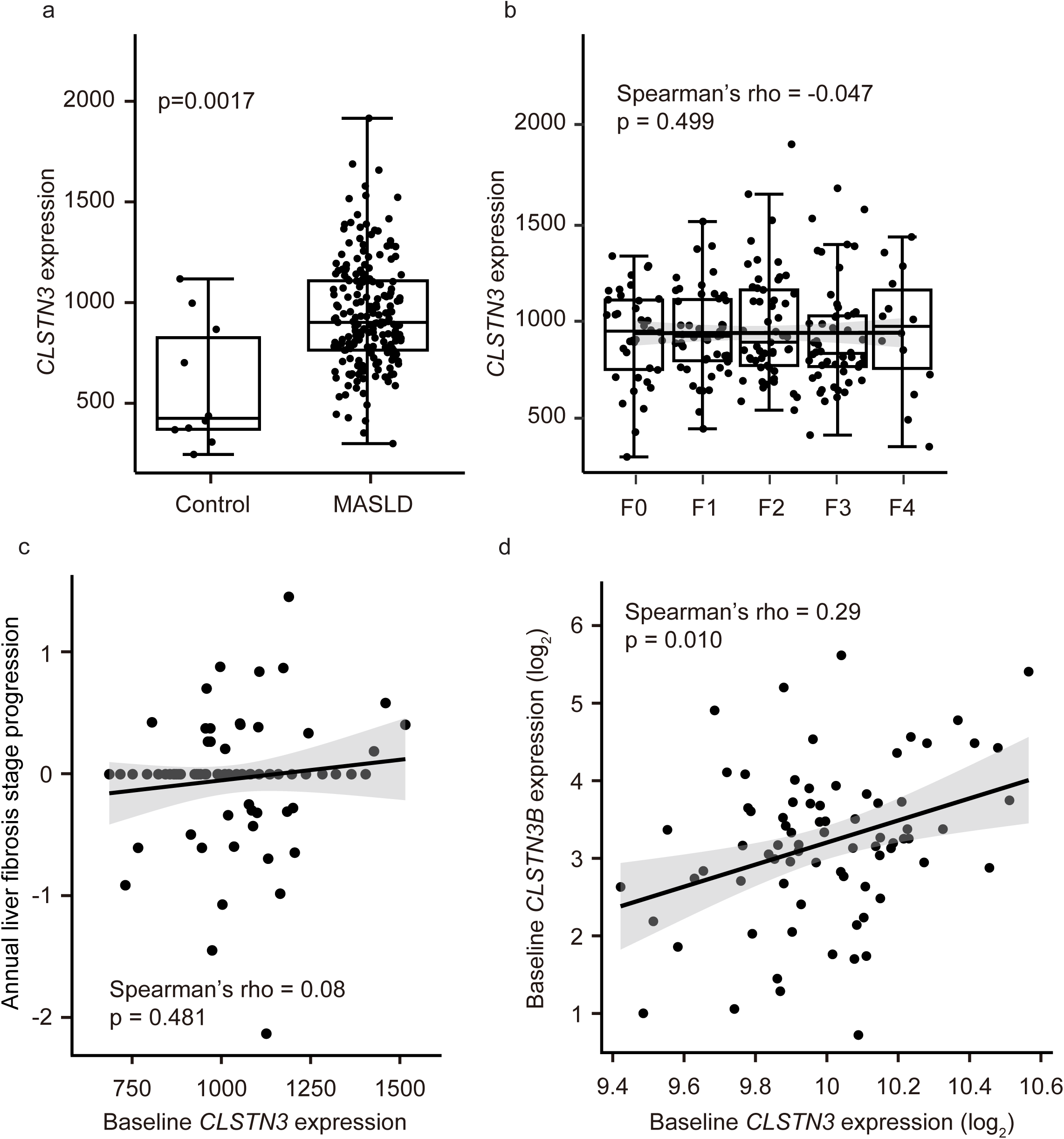
**a**, Hepatic *CLSTN3* expression levels in 206 European MASLD patients compared to 10 healthy obese controls. **b**, Hepatic *CLSTN3* expression levels by liver fibrosis (F) stage in 206 European MASLD patients. **c**, Correlation between baseline *CLSTN3* expression levels and time-adjusted (i.e., annual) progression of liver fibrosis stage in 77 Japanese MASLD patients. **d**, Correlation between baseline *CLSTN3* and *CLSTN3B* expression levels in 77 Japanese MASLD patients. Inter-group difference was assessed by Wilcoxon rank-sum test. Correlation was assessed by Spearman correlation test.

